# Structural and functional basis of PU.1-BAF interaction enables targeting of lineage-specific transcription

**DOI:** 10.1101/2025.08.01.667454

**Authors:** Dipti Sadalge, Jozlyn R. Clasman, Salih Topal, Derek Musser, Laura Von der Porten, Siying Wu, Samantha E. Schultz, Wesley Austin, David Huang, Ying-Duo Gao, Jeremy W. Setser, Marissa R. Martinez, Janna Kiselar, Stephen Hesler, Sydney J. Blevins, Bengi Turegun, Gavin M. Palowitch, Alexis Khalil, John L. Pulice, Mark R. Chance, Kevin J. Wilson, Marina Nelen, David L. Lahr, Gabriel J. Sandoval, Yunji W. Davenport, Steven F. Bellon, Asad M. Taherbhoy

## Abstract

Chromatin remodeling complexes like BAF finely regulate transcriptional programs by working in concert with transcription factors. However, evidence is lacking as to whether TFs interact directly with BAF and if so, what the mechanistic and structural principles governing these critical interactions are. Here, we establish direct engagement between a crucial and therapeutically relevant full-length human TF, PU.1 (*SPI1*), and BAF. Within this 1MDa+ complex, we precisely map the binding site of PU.1 to a YEATS-like domain on BAF60A and elucidate the structure of the PU.1-BAF60A complex. This work reveals that upon binding to BAF, a disordered region within the TF adopts a helical conformation, and that disruption of this functionally critical interface via knockdown abrogates the ability of PU.1 to rescue cell viability. To explore the druggability of TF-BAF protein-protein interactions (PPIs), we conducted a high-throughput screen that identified small molecules capable of disrupting the PU.1-BAF60A PPI by binding to BAF60A. Co-crystal structures reveal distinct compound binding modes that converge on a critical PU.1-BAF60A interaction hotspot. These findings define, for the first time, the structural interface between a human TF and a chromatin remodeling complex and establish a platform that enables the targeting of these interactions, a novel mechanism in cancer therapeutics.

## Introduction

Precise and timely regulation of global chromatin architecture is vital for the function of both mature and developing cells. To maintain a functional chromatin landscape, various classes of ATP-dependent chromatin remodeling complexes, such as BRG1/BRM-associated factors (BAF; also known as mammalian SWI/SNF), chromodomain helicase DNA-binding (CHD), and imitation switch (ISWI), are utilized by the cell to reposition nucleosomes and thereby modulate transcription^1^. The mSWI/SNF or BAF complex is a multi-subunit protein complex over 1MDa in size comprising 3 distinct classes, cBAF, PBAF, and ncBAF, with each class featuring unique subunits and different paralogs that define their identity and function^2–5^; notably, genes encoding these subunits have been found to be mutated in approximately 20% of all human cancers^6–8^. Examples include loss-of-function mutations, as observed in the tumor suppressor SMARCB1 subunit in malignant rhabdoid tumors^9^; ARID1A*-*deficiency or several mutations in numerous cancers, including liver, bladder, gastric, and ovarian clear cell carcinoma^10^; and fusion oncoproteins, such as the fusion of the BAF component SS18 to portions of SSX gene products in synovial sarcoma^11^. BAF can also drive oncogenesis through collaboration with oncogenic transcription factors (TFs), as exemplified by hijacking of BAF by fusion TFs like EWS-FLI1 in Ewing sarcoma^12^ and TMPRSS2-ERG in prostate cancer^13^.

Transcription factors bind to specific DNA sequences to regulate gene expression by modulating chromatin accessibility via the recruitment of chromatin regulators. Loss- and gain-of-function perturbations in TFs are implicated in various diseases including cancer^14,15^, neurodegenerative diseases^16,17^, and autoimmune disorders^18,19^. Many TFs are known to enable recruitment of BAF complexes to target sites to remodel chromatin accessibility and regulate gene expression, and these TFs depend on the integrity of BAF complex subunits^13^ and catalytic function^20^ for their activity. The importance of these TF-BAF complex collaborations has been validated for several TF families including ETS^12,13,21,22^, Forkhead^23^, AP-1^24^, POU2F3^25^, and nuclear hormone receptors^26–28^. While numerous studies have demonstrated that TFs work with BAF complexes to create genome-wide regulatory landscapes, it is not yet understood whether TFs interact directly with BAF, and if so, how.

Purine-rich box-1 (PU.1), an ETS family TF, is a master regulator of hematopoiesis, orchestrating lineage specification of both common lymphoid and granulocyte-macrophage progenitors and guiding terminal differentiation of monocyte-macrophages and B cells^21,29^. PU.1 is thought to act as both an oncogene and a tumor suppressor in AML. High expression of *SPI1* (PU.1) was associated with shorter overall survival in patients with intermediate cytogenetic risk AML (vs low *SPI1* expression)^30^ and reducing PU.1 levels decreased cell viability in AML^31^, making PU.1 an attractive therapeutic target. Therapeutic strategies targeting PU.1 include small-molecule inhibitors designed to disrupt the interaction of PU.1 with DNA^32^ or other transcription factors^33^ but concerns remain regarding the selectivity of these molecules. Given that PU.1 is a classical pioneer TF that can bind its sequence motif occluded on the nucleosome^34^ and promote the recruitment of chromatin remodeling complexes to alter chromatin accessibility^22,34–36^, the PU.1-BAF axis provides an ideal context for both mechanistic interrogation and therapeutic targeting of TF-BAF interactions in cancer.

Here, leveraging SMARCA2/4 inhibitors and knock-down of PU.1, we establish the interconnectedness of BAF and PU.1 in regulating transcriptional programs. Using an *in vitro* system with recombinant proteins, we provide, for the first time, an in-depth, structural understanding of how a human pioneer TF interacts with BAF. Finally, using robust high-throughput screening (HTS) we identify protein-protein interaction (PPI) inhibitors of the PU.1-BAF interaction, offering a new therapeutic strategy for targeting aberrant TF function.

## Results

### Cellular validation of the PU.1-BAF collaboration in AML

Pioneer TFs have been found to be key drivers of oncogenesis across multiple tumor types. One mechanism by which TFs drive tumor development and maintenance is through interactions with the BAF complex. Recently, small molecules inhibiting the ATPases SMARCA2 and SMARCA4 have been identified^37^, essentially inhibiting all BAF complex function in the cell. We used the PRISM platform^38,39^ to test the impact of dual SMARCA2/SMARCA4 ATPase inhibition with FHT-1015 (a potent SMARCA2/4 ATPase inhibitor^34,37^) across a wide range of human cancer cell lines (n=565). Dose-dependent growth inhibition was observed across numerous cell lines, with especially strong inhibition in several hematologic malignancies (leukemia and lymphoma) (Fig. 1a, Extended Data Fig. 1a). The effects in AML (Extended Data Fig. 1a) were of particular interest as this tumor type is thought to be driven by TF misregulation^22,40,41^.

**Fig. 1.**
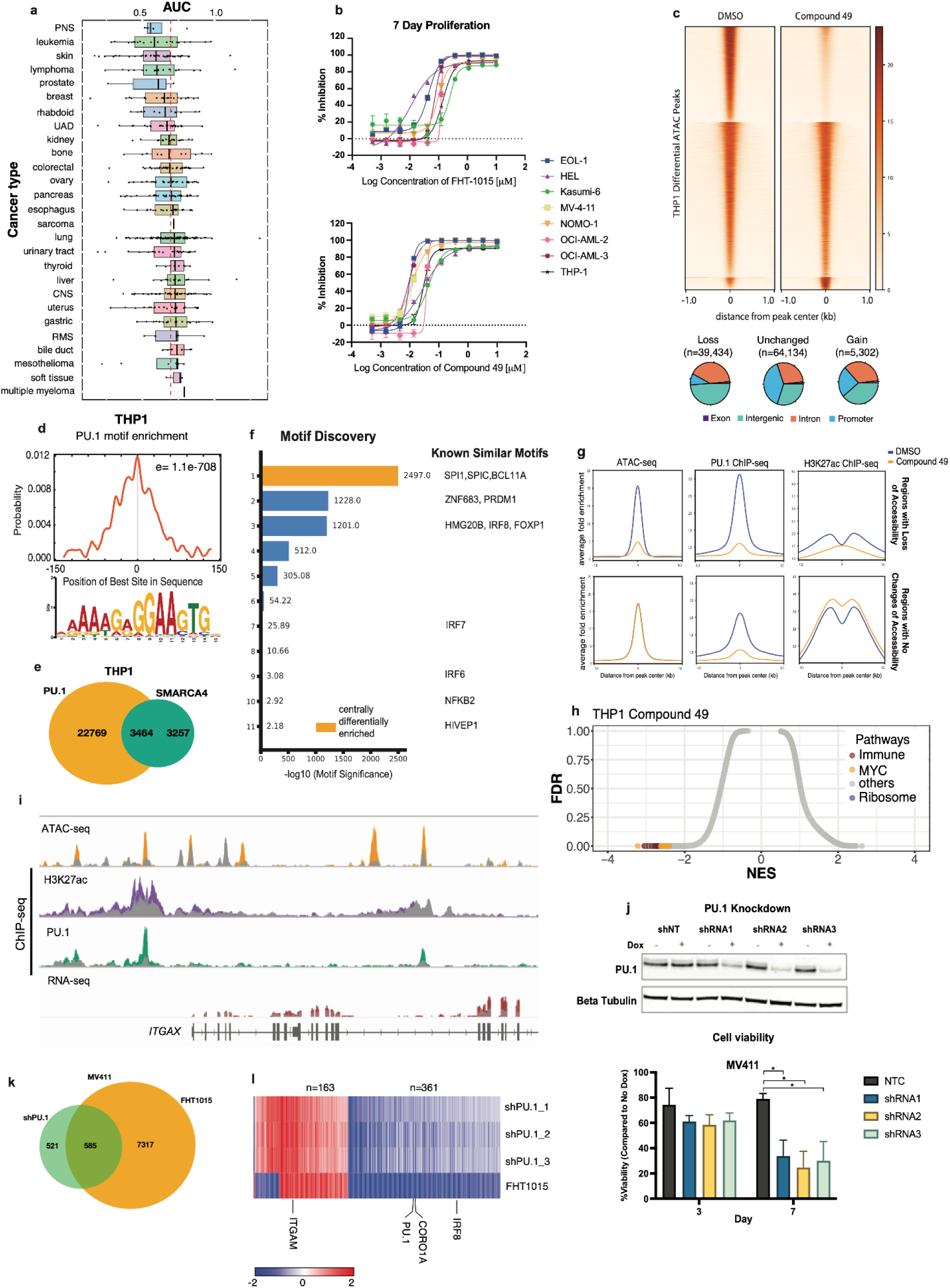
Cellular validation of SPI1 (PU.1)-BAF interplay in AML. **a,** Area Under the Curve ( AUC) values of a range of cancer cell lines (n=565) treated with FHT-1015 for 3 days. Each box displays the distribution of AUC values within each cancer type; vertical line represents median. Red dashed line represents the sensitivity threshold for FHT-1015 and values to the left of the dashed line indicate the sensitive cell lines. **b,** Percentage of proliferation inhibition in response to increasing concentrations of FHT-1015 (top graph) or Compound 49 (bottom graph) for 7 days in 8 AML cell lines. **c,** Tornado plot showing changes in chromatin accessibility (sorted by DMSO signal) in THP-1 AML cells in response to treatment with Compound 49 for 24h. Pie charts showing distribution of genomic features of ATAC regions by type of change in chromatin accessibility. **d,** Enrichment p-value and motif probability graph for SPI-1 as the most enriched motif at regions with loss of chromatin accessibility in THP-1 cells. **e,** Venn diagram showing overlap between PU.1-occupied vs SMARCA4-occupied genomic sites in THP-1 cells. **f,** Bar plot showing top motifs identified in SMARCA4-occupied regions. **g,** Anchor plots showing average signal for ATAC-seq (left), PU.1 ChIP-seq (middle), and H3K27ac ChIP-seq (right) over loss of accessibility regions (top) or unchanged regions (bottom) in THP-1 cells treated with Compound 49 (DMSO signal in blue, Compound 49 signal in orange). **h,** Gene set enrichment analyses (GSEA) of THP-1 cells treated with DMSO or Compound 49 for 24h. Differential pathway results were ranked and displayed as normalized enrichment scores (NES). **i,** Genome browser view of ATAC-seq, ChIP-seq of PU.1 and H3K27ac, and RNA-seq at the ITGAX locus in THP-1 cells treated with DMSO or Compound 49 for 24h. Gray signal on each track indicates Compound 49-treated samples. **j,** (Top) Western blot depicting PU.1 levels in MV-4-11 cells containing 3 doxycycline-inducible shPU.1 or a non-targeting control (shNT). Cells were treated with or without doxycycline for 48h. (Bottom) Cell viability normalized to “no doxycycline” control MV-4-11 cells in the presence of shRNAs targeting PU.1 (n=3 biological replicates). (*p < 0.05, unpaired t-test versus shNT). **k,** Venn diagram showing the overlap of differentially regulated genes (p_adj_ <0.05) in response to 48h shPU.1 knockdown or 24h FHT-1015 treatment. Adjusted *P* values were calculated using the Benjamini-Hochberg method. **l,** Heat map showing the overlapped genes (n=585) and highlighting key PU.1 target genes.

To evaluate the cell growth inhibition observed in the PRISM screen, 8 AML cell lines (EOL-1, HEL, Kasumi-6, MV-4-11, NOMO-1, OCI-AML-2, OCI-AML-3, and THP-1) were grown in the presence of increasing concentrations of either FHT-1015 or Compound 49, another SMARCA2/4 inhibitor with properties similar to that of FHT-1015 (Extended Data Fig. 1b and 37), for 3 or 7 days. Treatment with both compounds at nanomolar concentrations led to robust proliferation inhibition on day 3, and near total proliferation inhibition on day 7, confirming sensitivity to BAF inhibition in AML (Fig. 1b and Extended Data Fig.1c).

To understand the genomic features contributing to sensitivity to dual ATPase inhibition in AML, we performed a transposase-accessible chromatin with sequencing (ATAC-seq) assay to investigate changes in chromatin accessibility in the AML cell line THP-1, treated with Compound 49. Of the 108,870 regions detected through the chromatin accessibility assay, 36% (n=39,434) lost accessibility in response to Compound 49 (Fig. 1c). Most regions that lost accessibility were intronic or distal intergenic, which indicates that they are most likely enhancer regions (Fig. 1c). Five percent (n=5,302) of the regions detected by ATAC-seq gained chromatin accessibility in response to Compound 49 (Fig. 1c). Similar results were observed with FHT-1015 in the MV-4-11 AML cell line (Extended Data Fig. 1d). Motif analysis was performed on regions that had a loss of accessibility in both cell lines. Interestingly, this analysis found that the most enriched motif at these sites was *SPI1* (PU.1), suggesting that BAF ATPase activity might be critical for *SPI1* (PU.1) occupancy at these sites along the genome (Fig. 1d and Extended Data Fig.1e). PU.1 is important for the maintenance of hematopoietic stem cells (HSCs) and is critical for myeloid and lymphoid differentiation^42–44^. Interestingly, mice harboring loss of SMARCC1 (a core BAF complex subunit), displayed diminished PU.1 activity, suggesting the BAF complex may be required to establish PU.1 activity in cells^45^.

The identification of *SPI1* as the most commonly enriched motif in the genomic regions that became inaccessible upon BAF inhibition prompted us to investigate the relationship between *SPI1* (PU.1) and BAF at a genome-wide scale. To this end, we performed chromatin immunoprecipitation sequencing (ChIP-seq) to investigate the distribution of PU.1 across the genome, along with SMARCA4 (BAF) and H3K27ac (a marker for active enhancer regions), in THP-1 cells. We identified 26,233 genomic sites occupied by PU.1 and 6,721 sites occupied by SMARCA4 (Fig. 1e). Fifty-one percent (n=3464) of SMARCA4-occupied sites were co-occupied by PU.1, indicating an important relationship between PU.1 and the BAF complex. Additionally, the most commonly enriched motif across all SMARCA4-occupied sites was identified as *SPI1* (Fig. 1f). Considering the strong co-localization of PU.1 and SMARCA4, we investigated the effect of SMARCA2/4 inhibition on PU.1 occupancy at the open chromatin regions identified using ATAC-seq in THP-1 cells. As expected, Compound 49 treatment resulted in a substantial reduction in PU.1 occupancy at regions with loss of chromatin accessibility (n=39,434), and only a mild decrease in PU.1 occupancy at unchanged regions (n=64,314) (Fig. 1g). Next, we investigated the transcriptional effects in THP-1 cells exposed to 100 nM of Compound 49 for 24h. Gene set enrichment analysis (GSEA) showed strong downregulation of MYC-related pathways, along with ribosomal and immune system pathways, in response to Compound 49, suggesting that BAF activity is required for proliferative and developmental functions in AML cells (Fig. 1h). Notably, *ITGAX*, a critical monocyte differentiation gene and known PU.1 target, was strongly downregulated in response to treatment with Compound 49, correlating with a substantial reduction in chromatin accessibility along with loss in PU.1 and H3K27ac occupancies at the upstream region of the ITGAX locus (Fig. 1i). Taken together, these results highlight the functional relationship of PU.1 and BAF in AML.

Because *SPI1* (PU.1) was the most commonly enriched motif in chromatin regions that lost accessibility in response to BAF inhibition, and hematologic cell lines displayed substantial proliferation defects in response to BAF inhibition, we investigated whether PU.1 is directly required for leukemic cell survival. MV-4-11 cells were treated with 3 doxycycline-inducible shRNAs targeting PU.1, along with a non-targeting shRNA construct as a negative control. All 3 shRNAs targeting PU.1 significantly (p<0.05) reduced proliferation of MV-4-11 cells by day 7 (Fig. 1j). This data is in agreement with a publicly available CRISPR knockout dataset^46,47^ (Extended Data Fig. 1f). Next, transcriptional changes in response to PU.1 knockdown were investigated in MV-4-11 cells. Strikingly, more than half (n=585) of genes differentially regulated in response to shPU.1 for 48h were also differentially regulated in response to FHT-1015 treatment (Fig. 1k). Among these genes were known PU.1 target genes such as *IRF8*, *CORO1A*, and *SPI1* (PU.1) itself (Fig. 1l). The overlapping transcriptomic effects of TF knockdown and BAF inhibition reveals a strong genetic interplay between PU.1 and the BAF complex.

To further investigate whether the pioneering activity of PU.1 is mediated through interactions with BAF, we performed co-immunoprecipitation of V5-tagged PU.1 transiently overexpressed in HEK-293-T cells. The samples were immunoprecipitated using anti-V5 antibody and run via western blot. The results revealed that PU.1 can pull down the BAF complex (Extended Data Fig. 1g), providing additional evidence that PU.1 and BAF interact in a cellular context.

### *In vitro* confirmation of PU.1-BAF interaction

To understand whether the interaction between PU.1 and BAF is direct or indirect, we conducted pull-down experiments using recombinantly purified, maltose-binding protein (MBP)-tagged, full-length PU.1 (PU.1). Purified PU.1 and MBP only were immobilized on amylose beads and incubated with fully assembled cBAF complex. Upon running the samples using sodium dodecyl sulfate polyacrylamide gel electrophoresis (SDS-PAGE), bands corresponding to cBAF components were detected in samples pulled down using PU.1 but not in the samples pulled down by MBP alone, suggesting a direct interaction between PU.1 and cBAF without the involvement of other proteins (Fig. 2a, b). In subsequent pull-down experiments with either cBAF structural core or catalytic core (ATPase and ARP module), PU.1 pulled down the structural core but not the catalytic core, indicating that PU.1 interacts with a region within the structural core (Fig. 2b). SDS-PAGE analysis confirmed the purity and presence of subunits within cBAF, cBAF structural core, and cBAF catalytic core (Extended Data Fig. 2a). Size-exclusion chromatography coupled with multi-angle light scattering (SEC-MALS) further confirmed the size and integrity of each complex, and the elution profile was consistent with stable complexes at expected molecular weights, free of major aggregates (Extended Data Fig. 2b). To quantitate the interaction between PU.1 and cBAF, we employed analytical ultra centrifugation (AUC), in which the sedimentation coefficient (s) of cBAF was monitored at varying concentrations of PU.1. PU.1 bound to cBAF with K_D_ of 1.64 µM (representative value from 2 independent measurements) (Fig. 2c-e). We were thus able to establish a direct, mutually sufficient molecular interaction between PU.1 and cBAF.

**Fig. 2.**
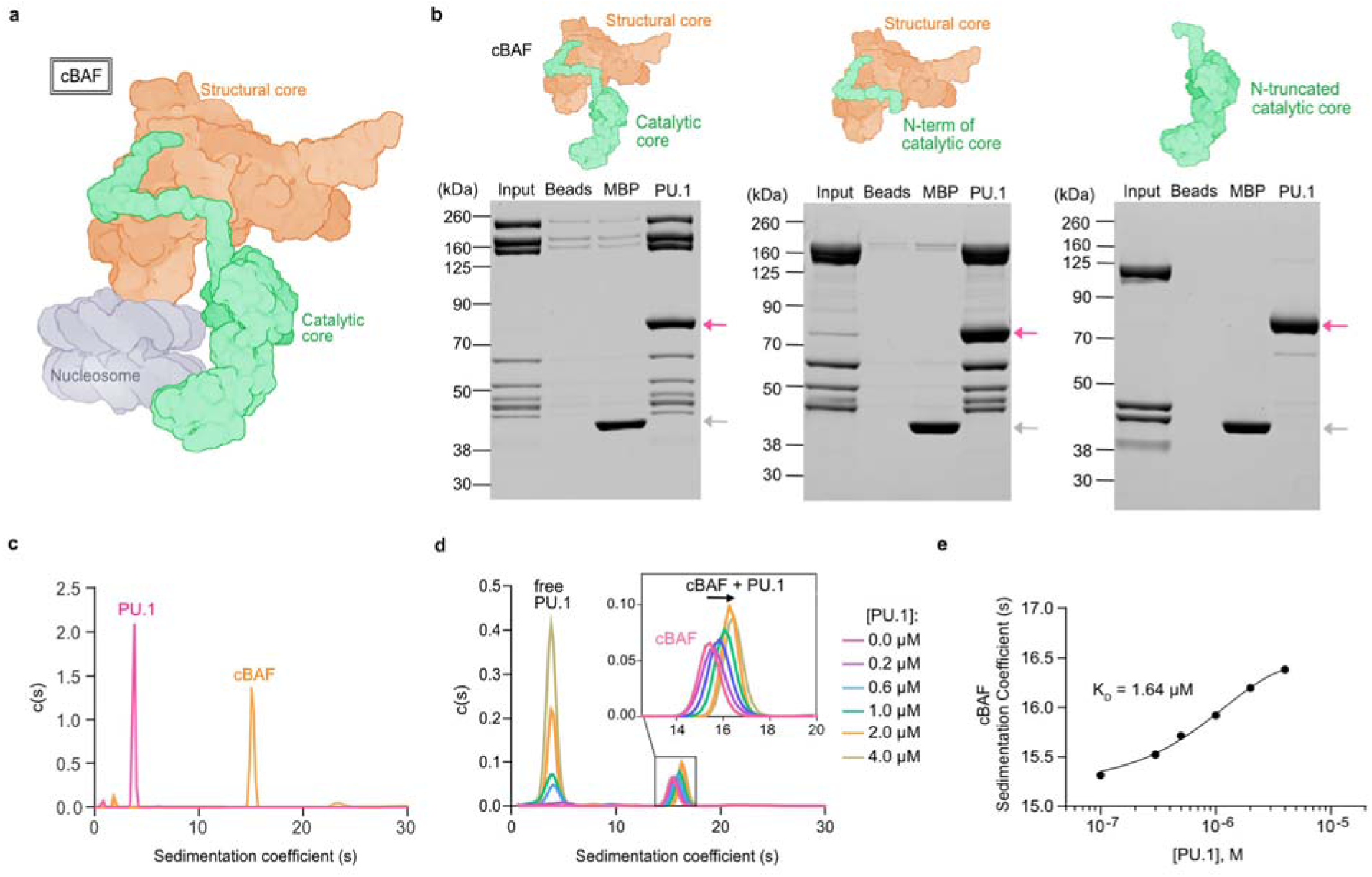
PU.1 interacts with the structural core of the BAF complex in vitro. **a,** Cartoon model representation of cBAF (based on PDB: 6LTJ)^56^. The structural core (orange), catalytic core (green), and nucleosome (gray) are labeled. **b,** Amylose pull-down assay demonstrating direct interaction of PU.1 with the cBAF complex and structural core and lack of direct interaction with cBAF catalytic core. MBP used as control. Pink and gray arrows indicate bands of recombinant PU.1 and MBP, respectively. Refer to Extended Data Fig. 2 for identity of input bands. **c,** Analytical ultracentrifugation (AUC) profiles showing sedimentation velocity [c(s)] of 8 µM PU.1 (pink) and 0.9 µM cBAF (orange). **d,** Sedimentation velocity [c(s)] analysis showing the sedimentation coefficient (s) of free PU.1 and dose-dependent shift in sedimentation coefficient of cBAF in complex with 0.2 to 4 µM PU.1. **e,** Sedimentation coefficient (s) of cBAF plotted against PU.1 concentrations. Data fit to 1-site binding equation using GraphPad Prism (version 10.0.2) to obtain K_D_.

### Mapping the PU.1 binding site to a specific region on cBAF

To map the specific PU.1 binding site on cBAF, we performed mass spectrometry-based protein footprinting that takes advantage of covalent chemistries to label solvent-accessible amino acid side chains on proteins. The presence of a ligand or binding partner shields the interaction surface from solvent, resulting in reduced labeling of amino acid side chains at the interface. Comparison of the modification rates of trypsin-digested peptides on the *apo* protein and those on protein in the presence of its binding partner provides insights into the interaction surface^48^. For our study, we used 2 complementary labeling chemistries that employ either 1-ethyl-3-(3-dimethylaminopropyl) carbodiimide (EDC) and glycine ethyl ester (GEE) to modify solvent-accessible carboxyl groups on glutamate (E) and aspartate (D)^48–50^ or CF3 radicals (•CF3) produced from a trifluoromethylation (TFM) reagent that targets a much broader range of residues^51,52^.

Upon PU.1 binding, mass spectrometry-based protein footprinting revealed protection (reduction in labelling) on 2 cBAF subunits: BAF60A and ARID1B. With EDC/GEE labeling, the 3 most protected peptides on BAF60A were 215-223 (site E217) and the contiguous peptides 239-246 (site E243) and 247-260 (sites D247 and D252) (Fig. 3a). CF3 labeling similarly showed protection on 2 contiguous BAF60A peptides, 247-260 (sites H254, W258, and H259) and 261-273 (site F270) (Fig. 3a). While CF3 did not identify any notable protected peptides on ARID1B, EDC/GEE labeling revealed 2 highly protected peptides, 1905-1911 and 2076-2094, both located in the disordered regions of ARID1B (Extended Data Table 1). The same ARID1B peptides were observed in our footprinting experiments with multiple other TFs (data not shown), suggesting that the ARID1B protected sites observed here are likely nonspecific or due to conformational changes induced by TF binding. Thus, we focused our attention on BAF60A, as these sites appeared unique to binding with PU.1.

**Fig. 3.**
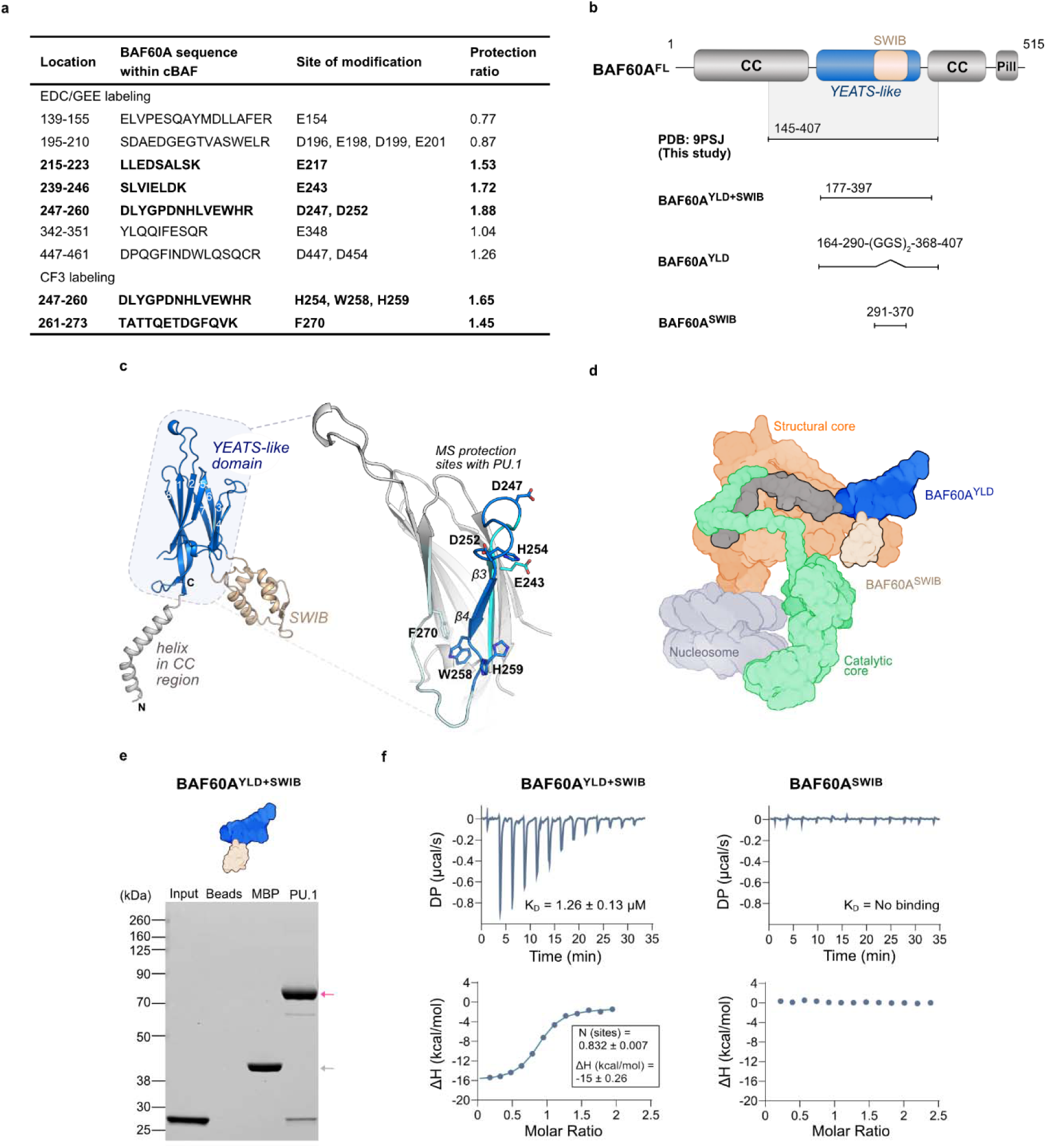
BAF60A YEATS-like domain in BAF is a mediator of PU.1 interaction. **a,** Table of MS footprinting data with protection ratios for BAF60A peptides following EDC/GEE and CF3 labeling of BAF reveals regions within BAF60A interacting with PU.1. The peptides with the highest degree of protection are bolded. **b,** Domain map of BAF60A (top) and BAF60A constructs designed for structural studies and to validate the region of BAF60A that interacts with PU.1. The domains of BAF60A are annotated as the coiled coil (CC, dark gray), YEATS-like (blue), SWIB (beige), and the pillar helix (Pill, dark gray) **c,** Structure of BAF60A apo highlighting the architecture of the N-terminal helix of CC region (gray), YEATS-like (blue) and SWIB domains (beige). The region corresponding to the top-ranking protections are colored in shades of blue, located between β3-β4 of the YEATS-like domain. The residue sites modified from MS footprinting are annotated. **d,** Model representation of cBAF (based on PDB: 6LTJ)^56^ integrating our BAF60A apo structure. The structural core (orange), BAF60A^YLD^ (blue), BAF60A^SWIB^ (beige), catalytic core (green), and nucleosome (gray) are labeled. **e,** Amylose pull-down demonstrating direct interaction between PU.1 and BAF60A^YLD+SWIB^. Pink and gray arrows indicate bands of recombinant PU.1 and MBP respectively. **f,** ITC data for PU.1 binding to BAF60A^YLD+SWIB^ or BAF60A^SWIB^. Upper panels show baseline-adjusted raw power data from titrations, and lower panels display 1-site binding kinetics fits, obtained using MicroCal PEAQ-ITC Analysis Software (version 1.41).

**Table 1.**
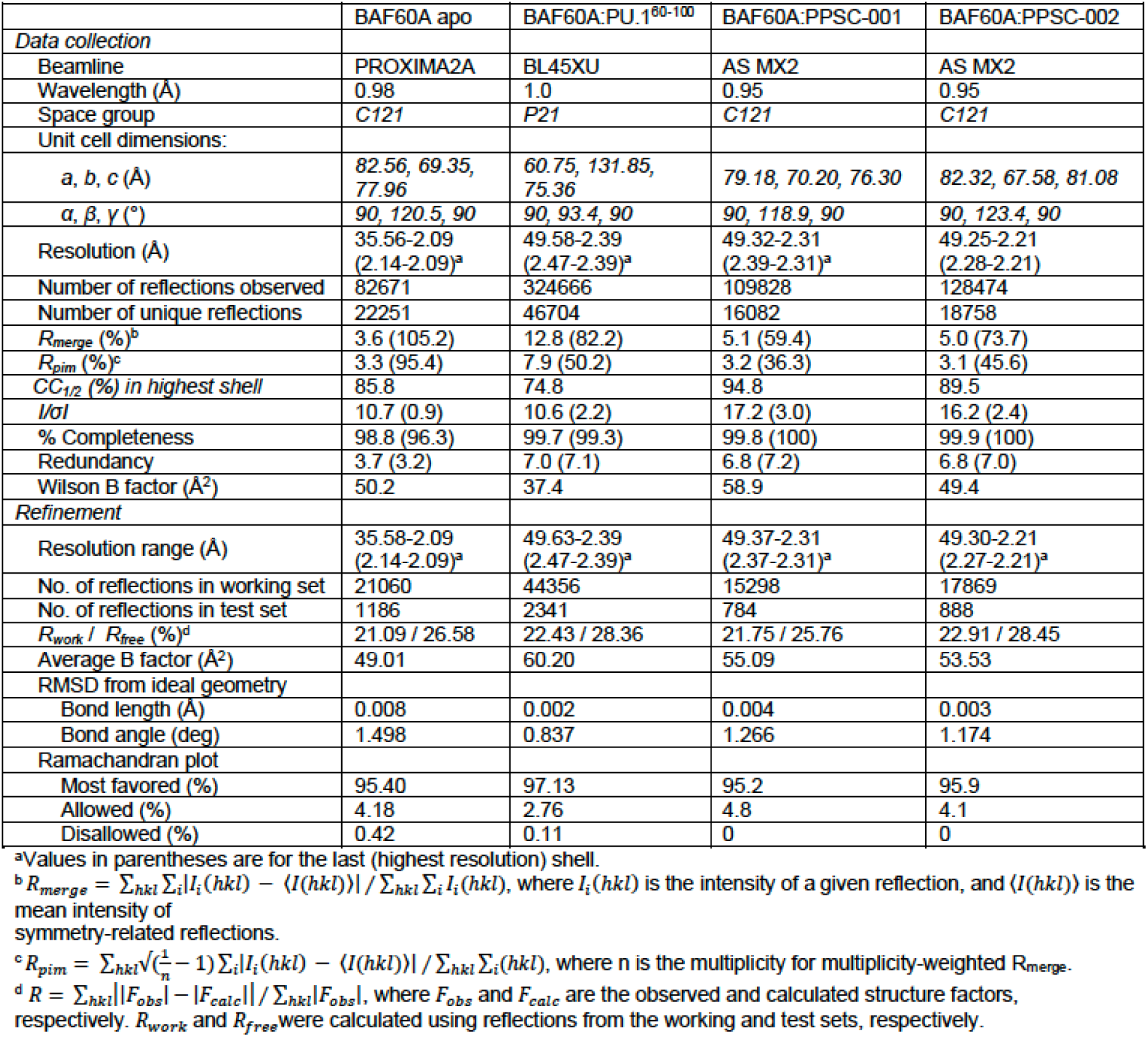
Cata collection and refinement statistics.

All protected sites on BAF60A were located within the YEATS-like domain (YLD) of BAF60A (BAF60A^YLD^). This domain was first structurally characterized in pBAF^53^ (Protein Data Bank [PDB] code: 7Y8R) via cryo-electron microscopy at 4.4 Å, along with its yeast counterpart Snf12^54,55^, but was unresolved in human cBAF structures^56,57^. Since all prior structures were solved within a larger complex and with modest resolution, we determined for the first time the isolated X-ray crystal structure of BAF60A at 2.1 Å resolution to better visualize its protected sites upon TF binding (Table 1). BAF60A crystallized in the C2 space group with 1 copy in the asymmetric unit and contained residues spanning 144-407 of full-length protein, which includes a segment of the first helix of the coiled coil (CC) at the N-terminus and a well-defined YLD topologically linked to the SWI/SNF complex B (SWIB) domain^58^ (Fig. 3b, c). We will refer to this region of BAF60A as BAF60A^YLD+SWIB^. The BAF60A^YLD^ domain adopts an 8-stranded β-sandwich fold with 2 parallel 4-stranded β-sheets, as seen in YEATS domain family members such as ENL (*MLLT1*)^59^ and AF-9 (*MLLT3*)^60^. Structurally, BAF60A^YLD^ aligns well with canonical ENL^YEATS^, with a root mean square deviation (RMSD) of 2.19 Å (Extended Data Fig. 6a). The sites on BAF60A with the highest protection ratios, based on mass spectrometry-based protein footprinting, are all located on the same face of BAF60A^YLD^ between β3-β4 (Fig. 3c), with the exception of the protected area at peptide 215-223 (site E217), which is likely dynamic and was unresolved in our structure. To visualize the location and orientation of BAF60A^YLD^ within cBAF, we overlaid our structure with that of cBAF (PDB: 6LTJ)^56^ using BAF60A^SWIB^. Our structure overlays well with the resolved portion of BAF60A in the structural core of human cBAF, where the CC and BAF60A^SWIB^ both contact regions of BAF170 heterodimer and the armadillo repeats (ARM) of ARID1A (Extended Data Fig. 2c). BAF60A^YLD^ remains completely solvent-exposed and can therefore allow for the interaction of a protein partner like PU.1. A cartoon representation of how BAF60A integrates in cBAF is depicted in Fig. 3d.

In a pull-down assay, PU.1 was able to pull down the isolated BAF60A^YLD+SWIB^, confirming that the protection observed for BAF60A in the footprinting assay was due to a direct interaction between PU.1 and BAF60A, rather than conformational changes (Fig. 3e). To measure the binding affinity of PU.1 to BAF60A, we performed isothermal titration calorimetry (ITC). Titration of BAF60A^YLD+SWIB^ into PU.1 resulted in a K_D_ of 1.26 µM, with the molar ratio (n) supporting a 1:1 stoichiometry (Fig. 3f). This measurement was consistent with the dissociation constant of 1.64 µM obtained from AUC experiments with PU.1 binding to cBAF. BAF60A^SWIB^ alone did not bind to PU.1, suggesting that BAF60A^YLD^ is critical for this interaction (Fig. 3f). We thus mapped and quantitatively verified the BAF60A^YLD+SWIB^ domain as critical for the interaction between PU.1 and cBAF.

### A disordered region within PU.1 binds to BAF60A

Akin to the architecture of other TFs, PU.1 structurally features predominantly intrinsically disordered regions (IDRs) apart from its DNA-binding domain (DBD), which is highly ordered spanning residues 165-260 near the C-terminus (Fig. 4a). Proximal to the DBD, towards the N-terminus, is a negatively charged PEST domain (residues 117-165) that helps modulate the activity of PU.1 by regulating the dynamics of PU.1 homodimerization and therefore the DNA binding affinity of the DBD^61^. The N-terminal region contains an acidic transactivation domain (TAD) (residues 1-80) that is known to be involved in transcriptional activation and mediates interactions with other cofactors and protein partners^62^. Between the TAD and PEST domain is a glutamine-rich region (Q-rich) whose function is not well defined but may be similar to that of the TAD domain (Fig. 4b).

**Fig. 4.**
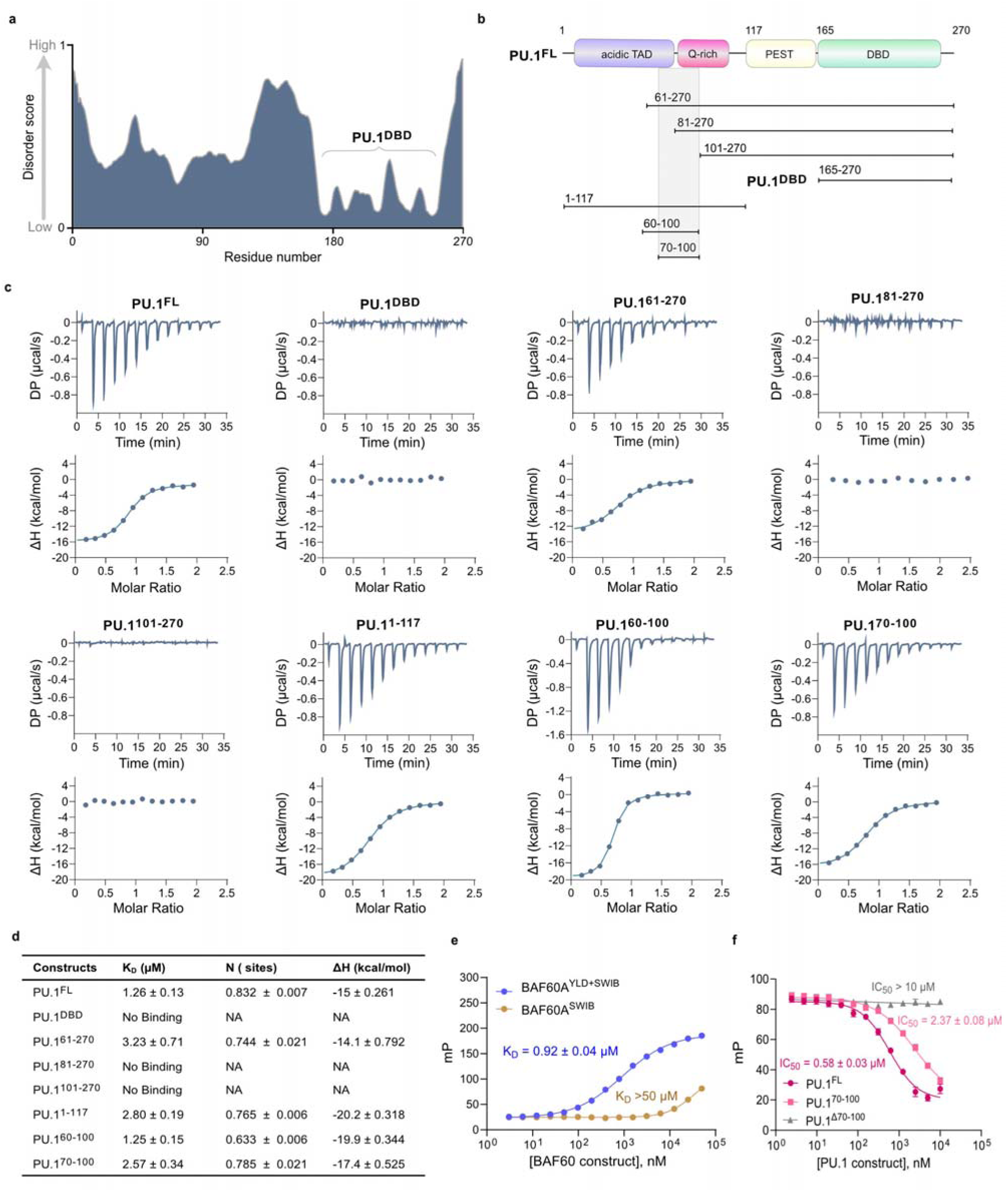
PU.1 associates with BAF60A YEATS-like domain via its N-terminal disordered region. **a,** Predicted disorder tendency of PU.1 on a scale of 0 (low tendency) to 1 (high tendency), calculated by Protein DisOrder prediction System (PrDOS). Position of the PU.1 DNA-binding domain (DBD) is indicated. **b,** Domain map of PU.1 (top) and PU.1 constructs designed to delineate the region of PU.1 that interacts with BAF60A. **c,** ITC data for the binding of the indicated PU.1 constructs to BAF60A^YLD+SWIB^. Upper panels show baseline-adjusted raw power data from titrations, and lower panels display 1-site binding kinetic fits obtained using MicroCal PEAQ-ITC Analysis Software (version 1.41). **d,** Summary of binding affinities and thermodynamic parameters derived from ITC experiments in **c**. The data include dissociation constants (K_D_), enthalpy changes (ΔH), and stoichiometry (N) for each PU.1 construct interacting with the BAF60A^YLD+SWIB^. **e,** FP titration curves for binding of PU.1^60–100^^TMR^ to BAF60A^YLD+SWIB^ (blue) and BAF60A^SWIB^ (beige) **f,** FP competition curves for PU.1^FL^ (pink circles), PU.1^70–100^ (pink squares), and PU.1^Δ70–100^ (gray) competing with PU.1^60–100^^TMR^ for binding to BAF60A^YLD+SWIB^. Data analyzed in GraphPad Prism (version 10.0.2) using a 4-parameter dose-response curve to obtain K_D_ ± SEM (panel **e**) and IC_50_ ± SEM (panel **f**) values. Error bars denote standard deviation from triplicates.

Previous studies indicated that the N-terminal disordered region of PU.1 is crucial for recruiting the BAF complex and accessing closed sites within compacted chromatin^36^. To identify the specific region that interacts with BAF60A, we made a series of N- and C-terminal PU.1 truncations (Fig. 4b) and tested their binding to BAF60A^YLD+SWIB^ by ITC (Fig. 4c, d). PU.1^DBD^ did not bind to BAF60A^YLD+SWIB^, indicating the region of PU.1 that interacts with BAF60A is outside the DBD. PU.1^61–270^ and PU.1^1–117^ bound to BAF60A^YLD+SWIB^ with a K_D_ of 3.23 µM and 2.80 µM, respectively, while PU.1^81–270^ and PU.1^101–270^ failed to bind BAF60A^YLD+SWIB^ at the concentrations tested (Fig. 4c, d). Based on the results from these truncations, we speculated that PU.1^60–100^ is the core region that associates with BAF60A. Indeed, PU.1 peptides PU.1^60–100^ and PU.1^70–100^ bound to BAF60A^YLD+SWIB^ with a K_D_ of 1.25 µM and 2.57 µM, respectively, comparable to the affinity of full-length PU.1 (PU.1^FL^) for BAF60A^YLD+SWIB^ (Fig. 4c, d). Additionally, BAF60A^YLD+SWIB^ binding to PU.1^FL^ and PU.1^70–100^ produced almost identical thermodynamic signatures, indicating that residues 70-100 were sufficient to recapitulate the binding of PU.1^FL^ (Extended Data Fig. 3a). Our ITC results were further confirmed with pull-down assays, in which PU.1^61–270^, PU.1^1–117^, and PU.1^70–100^ were able to pull down BAF60A^YLD+SWIB^ but PU.1^81–270^ and PU.1^101–270^ were unable to do so (Extended Data Fig. 3b). Having narrowed down the minimum necessary region for the interaction, we confirmed the affinity of this PU.1 region for BAF60A^YLD+SWIB^ in a fluorescence polarization (FP) direct binding assay that employed the PU.1 peptide encompassing residues 60-100 with an S60C mutation introduced to enable maleimide-based labelling with BODIPY TMR (PU.1^60–100^^TMR^). PU.1^60–100^^TMR^ bound to BAF60A^YLD+SWIB^ with K_D_ of 0.92 µM and showed no signs of interaction with BAF60A^SWIB^ (K_D_>50 µM; Fig. 4e). To verify this result, competition-based FP experiments were performed, resulting in highly comparable IC_50_ values for PU.1^FL^ (IC_50_: 0.58 µM) and PU.1^70–100^ (IC_50_: 2.37 µM) (Fig. 4f). S60C did not affect the binding of PU.1 to BAF60A^YLD+SWIB^, as demonstrated by an FP competition assay (Extended Data Fig. 3c). A PU.1 deletion construct in which the BAF60A interaction region (residues 70-100) was removed (PU.1^Δ70–100^) completely abolished binding to BAF60A^YLD+SWIB^ (Fig. 4f). Collectively, these results demonstrate that PU.1^70–100^ recapitulates the binding capability of PU.1^FL^.

### Structural features of the BAF60A:PU.1 complex

To elucidate the structural basis for PU.1 recognition by BAF60A, we determined the co-crystal structure of BAF60A^YLD+SWIB^ bound to PU.1^60–100^ at 2.5 Å resolution (Table 1). We attempted crystallization trials with both PU.1^60–100^ and PU.1^70–100^ but only PU.1^60–100^ was successful. The BAF60A^YLD+SWIB^: PU.1^60–100^ complex crystallized in a P2_1_ space group with 2 complexes oriented as an asymmetrical dimer on the 2-fold axis, resulting in 4 complexes in the asymmetric unit. The distinct face of BAF60A that was protected in the MS-based footprinting experiments, located between β3-β4, is engaged by an approximately 20-residue llJ-helix adopted by PU.1^60–100^ (Fig. 5a). Thus, PU.1^60–100^ likely transitions to a more ordered state upon binding to BAF60A, as is typical of classical IDRs.

**Fig. 5.**
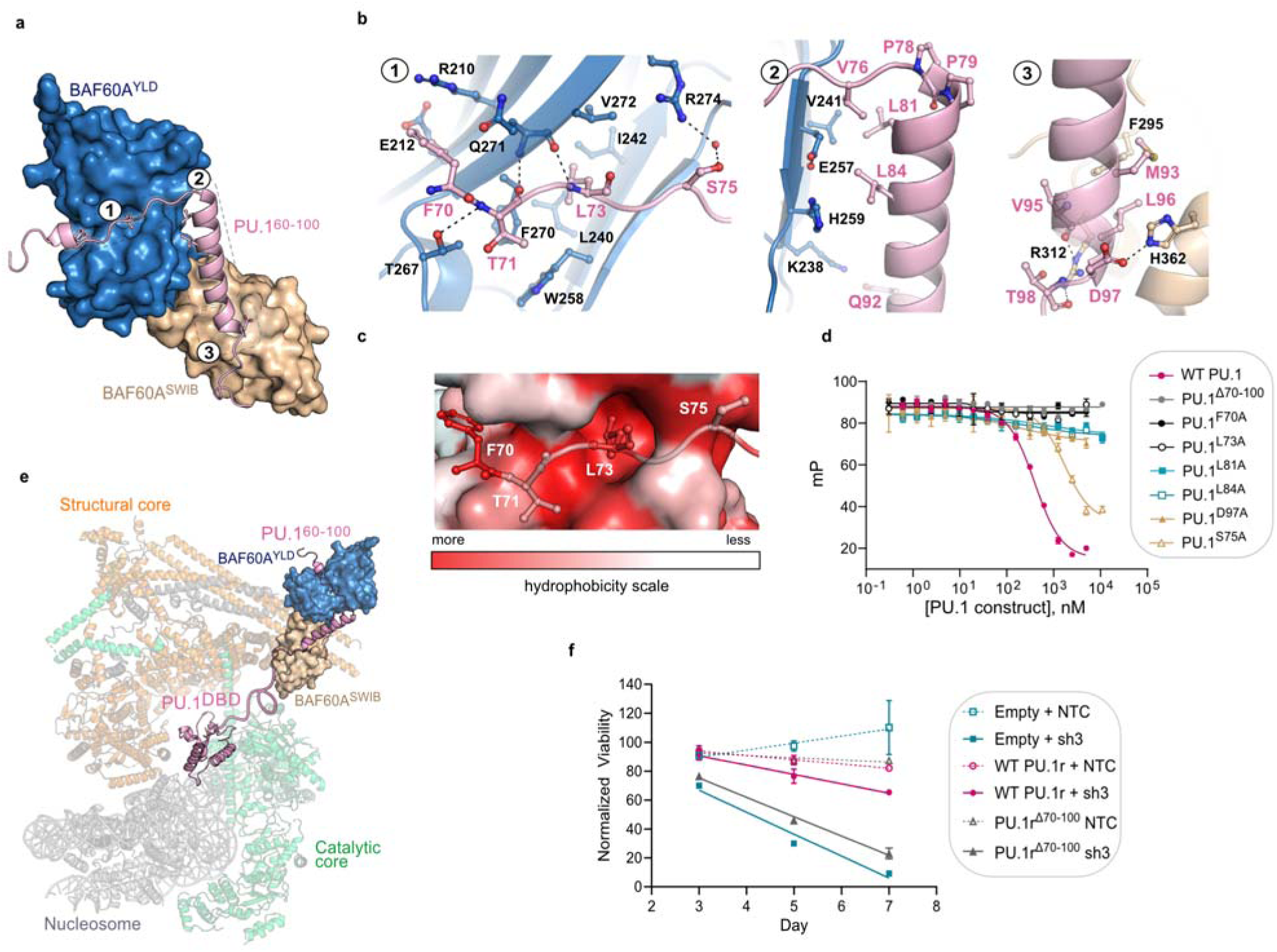
BAF60A interacts with PU.1 at 3 distinct binding pockets with both hydrophobic and hydrogen-bonding interactions. **a,** Structure of BAF60A^YLD+SWIB^: PU.1^60–100^ peptide complex. **b,** View of detailed protein-protein interactions at 3 locations designated 1-3 (also indicated in **a**). Region 1: Loop interactions involving F70 and L73 of PU.1. Region 2: PU.1 helix interactions involving hydrophobic contacts at L81 and L84 of PU.1. Region 3: PU.1 helix interactions involving D97 of PU.1 with BAF60A^SWIB^. **c,** Surface representation of Region 1 shaded to indicate hydrophobicity (low [white] to high [red]), highlighting important hydrophobic pockets for interaction. **d,** FP competition curves for PU.1 mutants competing with PU.1^60–100^^TMR^ for binding to BAF60A^YLD+SWIB^, compared to WT PU.1. Errors denote standard deviation from triplicates. IC_50_ values are provided in Extended Data Table 2. FP data analyzed in GraphPad Prism (version 10.0.2) using a 4-parameter dose-response curve to obtain IC50 ± SEM values. **e,** Model of PU.1 binding to cBAF using available structural information. Our structure and PU.1^DBD^ (PDB: 8EVH)^63^ was aligned to cBAF (PDB code: 6LTJ)^56^. PU.1^101–164^ (line between 60-100 and DBD) is approximately ∼60 Å. **f,** Cell viability of MV-4-11 cells containing empty vector, V5-PU.1r, or V5-PU.1r^Δ^^70–100^ with either shPU.1#3 or shNTC (+/- doxycycline). Cell viability was measured with Cell Titer Glo and was normalized to controls not incubated with doxycycline (n=2, each with 3 technical replicates).

In total, BAF60A engages with PU.1^60–100^ at 3 distinct regions covering a surface area of approximately 1500 Å^2^. Region 1 involves interactions at the N-terminal residue 70-78 loop of PU.1. Here, F70 sits in a groove between E212 and R210 and L73 inserts into a highly hydrophobic pocket within the BAF60A^YLD^ hydrophobic core, which is approximately 4 Å from MS footprinting protected sites F270 and W258 (Region 1, Fig. 5b). Region 2 encompasses van der Waals (VDWs) made by L81 and L84 within the llJ-helix of PU.1 at residues 80-97, where L84 interacts with protected site H259 on BAF60A^YLD^ (Region 2, Fig. 5b). Region 3 involves additional hydrophobic contacts and a notable hydrogen bond between D97 of PU.1 and H362 of BAF60A^SWIB^ (Region 3, Fig. 5b). Taken together, these results indicate that the IDR of PU.1^60–100^ changes form to enable 3 key points of contact via the formation of the llJ-helix to, first, ensure key residues on either end of the helix are able to engage BAF60A^YLD^ and BAF60A^SWIB^ (Region 2 and 3) and, second, direct the placement of Region 1 loop N-terminal to the helix via a sharp approximately 85° turn facilitated by 2 proline residues (P78 and P79) at the junction of the loop-to-helix transition (Region 2, Fig. 5b). This structural rearrangement allows the ideal trajectory for the 70-78 loop region and proper contact of F70 and L73 for engaging BAF60A^YLD^ (Region 1, Fig. 5b). To our knowledge, this binding mode appears completely novel, as a DALI search of the PDB did not identify any IDRs with structural similarity to the PU.1^60–100^ conformation.

To identify the critical binding residues within each region, a series of 6 single-point mutations were introduced into PU.1^FL^ and tested via competition-based FP. Three PU.1 point mutations were tested in Region 1, within the 70-78 residue loop (F70A, L73A, S75A). Two mutations were tested in the PU.1 helix at Region 2 (L81A, L84A) and 1 mutation was tested in the area of Region 3 contacting BAF60A^SWIB^ (D97A). Overall, the results corroborate our structural observations. Within the 70-78 loop of PU.1, F70A and L73A completely abolished binding to BAF60A^YLD+SWIB^. Substitution of an alanine at F70 removes VDWs interactions with the aliphatic chains of E212 and R210 of BAF60A^YLD^ and a potential cation-pi interaction with R210. L73 is crucial for interaction with BAF60A, as its side chain inserts into the BAF60A^YLD^ hydrophobic core, comprising of residues L240, I242, W258, F270, and V272. The S75A substitution had less of an impact, likely because it is involved in a water-mediated hydrogen bond. In the PU.1 helix at Region 2, L81 and L84 mutations disrupt VDWs contacts with V241 for L81A and E257 and with H259 for L84A. Finally, in Region 3, mutation of D97 disrupts the hydrogen bond with H362 of BAF60A^SWIB^, drastically reducing PU.1 binding.

To further probe the importance of BAF60A^SWIB^ in the PU.1 interaction, we tested BAF60A^YLD^ binding to PU.1^60–100^^TMR^ in an FP direct binding assay and found that removal of BAF60A^SWIB^ resulted in a >20-fold reduction in PU.1 binding affinity (Extended Data Fig. 4a, d), suggesting that BAF60A binding to PU.1 requires both BAF60A^YLD^ and BAF60A^SWIB^. We hypothesize that the helix of PU.1 acts as a scaffolding component to position the 70-78 loop for optimal binding. Therefore, disruption of the helix by removing BAF60^SWIB^ would also disturb the placement of the PU.1 70-78 loop within BAF60A^YLD^. We were unable to test the interaction from the BAF60A side, since mutating the relevant residues disrupted the hydrophobic core critical for protein stability. These results support a multifaceted interaction, with critical hot spots in each of the 3 regions of PU.1^60–100^ that contact BAF60A.

Previous work by Lian et al and He et al established the Cryo-EM structures of PU.1^DBD^- nucleosome (PDB: 8EVH)^63^ and BAF-nucleosome (PDB: 6LTJ)^56^. We combined these 2 structures by overlaying their common component, the nucleosome, revealing how BAF and a TF may interact simultaneously with a nucleosome. We next incorporated our structure by overlaying BAF60A^SWIB^ in our structure with the BAF60A^SWIB^ region of 6LTJ, since the BAF60A^YLD^ is not resolved in 6LTJ. This allowed us to propose a structural model with a 3-way interaction between a TF, BAF, and a nucleosome, highlighting a potential mechanism by which a TF could aid BAF recruitment to chromatin (Fig. 5e). In this model, the TF DBD binds the nucleosome, while the TF IDR acts as a molecular tether, binding BAF directly to help promote BAF interactions with the nucleosome. The distance between the C-terminus of PU.1^60–00^ and the N-terminus of PU.1^DBD^ can comfortably be spanned by the 64 residues of PU.1 that are also predicted to be disordered. One deficit of this model is that it lacks several regions of the BAF complex, which could impact how the TF-BAF-nucleosome complex assembles. Additional structural work will be needed to fully elucidate this 3-component complex.

There are 3 mutually exclusive paralogs of BAF60 that exhibit partially redundant functions yet display distinct tissue-specific expression patterns and distribution profiles^5^. Since the region of BAF60A that is involved in interacting with PU.1 is highly conserved within BAF60A, BAF60B, and BAF60C (Extended Data Fig. 4b), we expected PU.1 to maintain its binding affinity across all 3 paralogs. This was confirmed by an FP direct binding assay (Extended Data Fig. 4c, d). Therefore, it is likely the BAF60B and BAF60C paralogs interact with PU.1 via the same interface as BAF60A.

To determine whether the PU.1-BAF interaction is critical for cellular function, we engineered a PU.1 overexpression construct resistant to shRNA knock-down of PU.1 (V5-PU.1r). We additionally created a version of PU.1r that lacked residues 70-100 (V5-PU.1r^Δ70–100^). These constructs were then expressed in MV-4-11 cells previously infected with shPU.1 (+/- doxycycline). PU.1 expression was determined by Western blot analysis of cell lysates (Extended Data Fig. 4e). To probe whether V5-PU.1r (with or without residues 70-100) can rescue the cell viability phenotype observed with loss of PU.1 (Fig. 1g), a Cell Titer Glo assay was used to measure viability of cells expressing either V5-PU.1r or V5-PU.1r^Δ70–100^ (+/- doxycycline). Notably, wild-type (WT) V5-PU.1r was able to rescue the loss of cell viability caused by shRNA knockdown; however, V5-PU.1r^Δ70–100^was unable to rescue the cells from WT PU.1 loss (Fig. 5g). These data suggest that the PU.1-BAF interaction mediated by residues 70-100 of PU.1 serves critical cellular functions.

### Identification and characterization of novel PPI inhibitors of PU.1-BAF60A

Given the functional importance of the PU.1-BAF60A PPI, we next searched for a small molecule that could disrupt this interaction. Because BAF60A^YLD^ and ENL^YEATS^ share a common structural fold, we began by testing whether compound 94, an established ENL^YEATS^ binder, would also bind to BAF60A^YLD+SWIB^. (Crotonylated H3K27 [H3K27Cr] peptide, an ENL^YEATS^ binder that shares the same binding site as compound 94, served as a control.) Neither the compound nor the peptide bound to BAF60A^YLD+SWIB^ (Extended Data Fig. 6c-e). This finding was not surprising, given not only that PU.1 binds a site distal from the binding site of compound 94 and H3K27Cr, but also because BAF60A^YLD^ lacks the sequence conservation in the acylated lysine binding pocket possessed by other YEATS members (Extended Data Fig. 6a,b)^60,65^

In the absence of available small molecule disruptors of the BAF60A-PU.1 PPI, we used our previously described FP assay to screen for potential inhibitors. A high-throughput screening (HTS) campaign was conducted in a 1536-well format on a diverse compound library from Pivot Park Screening Centre (303,871 compounds, 30 µM compound conc., 1% dimethyl sulfoxide [DMSO]). The screen leveraged our robust FP assay with Z >0.7 and stable mP values for neutral and inhibition controls throughout the screen (Extended Data Fig. 5). A total of 155 inhibitors were identified using the 3-standard deviation cutoff of 18% (Fig. 6a). After a second round of validation in dose response and an orthogonal SPR (surface plasmon resonance) assay, 2 inhibitors, PPSC-001 and PPSC-002, were chosen for further characterization. PPSC-001 is a fused 1,4-dihydropyridine bearing a bromophenyl substituent at the 1 position, while PPSC-002 is a methylbenzoate substituted dihydro-dioxinobenzimidazole. PPSC-001 inhibited PU.1-BAF60A^YLD+SWIB^ interaction with an IC_50_ of 9.13 µM in the FP assay and bound to BAF60A^YLD+SWIB^ with a comparable K_D_ of 8.89 µM in the SPR assay (Fig. 6b, c). Similarly, PPSC-002 inhibited the PU.1-BAF60A^YLD+SWIB^ interaction with an IC_50_ of 33 µM in the FP assay and bound to BAF60A^YLD+SWIB^ with a comparable K_D_ of 22.63 µM in the SPR assay (Fig. 6b, c and Extended Data Fig. 6d). To probe the selectivity of these hits, we tested their binding to ENL^YEATS^ by SPR; no off-target binding was observed (Extended Data Fig. 6c). Thus, we determined that both PPSC-001 and PPSC-002 disrupted the PU.1-BAF60A interaction by specifically binding to BAF60A.

**Fig. 6.**
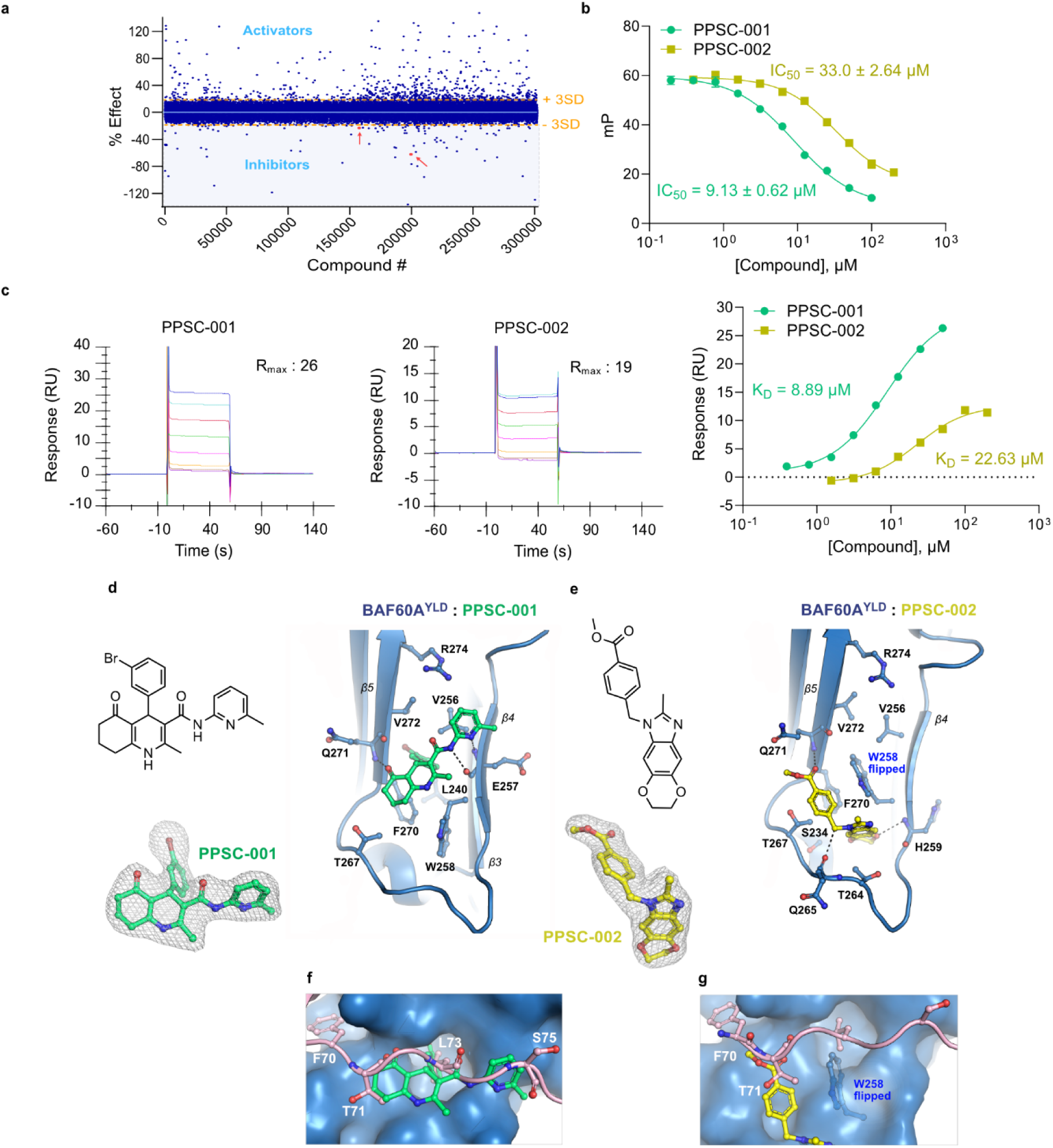
Hit identification, validation and structural characterization of BAF60A:PU.1 inhibitors. **a,** Overview of primary HTS. Hits (inhibitors) defined by compounds that exhibit percent inhibition of greater than 3 times the standard deviation (3SD=18%). Red arrows indicate the hits PPSC-001 and PPSC-002. Data analysis and visualizations were completed using GraphPad Prism (version 10.0.2). **b,** FP competition curves of PPSC-001 (green circles) and PPSC-002 (yellow squares) competing with PU.1^60–100^^TMR^ for binding to BAF60A^YLD+SWIB^. Error bars denote standard deviation from triplicates. Data analyzed in GraphPad Prism (version 10.0.2) using a 4-parameter dose-response curve to obtain IC_50_ ± SEM values. **c,** SPR binding curves and steady-state affinity measurements of PPSC-001 and PPSC-002 binding to BAF60A^YLD+SWIB^ in SPR (left and center). Expected saturation response (R_max_) for 1:1 stoichiometry is indicated. Equilibrium fits generated in GraphPad Prism (version 10.0.2) using a 4-parameter dose-response curve, to obtain K_D_ values (right). **d,e** Co-crystal structures of BAF60A^YLD+SWIB^ bound to **d,** PPSC-001 (green) and **e,** PPSC-002 (yellow) outlining detailed hydrophobic and peptide backbone interactions between β4-β5 of BAF60A^YLD^. The chemical structures and 2F_o_-F_c_ density maps of each compound are shown contoured to 1σ. **f,g** Extent of overlap between binding sites of **f,** PPSC-001 and **g,** PPSC-002 and PU.1 (pink cartoon) in BAF60A^YLD^ (surface representation from BAF60A:PU.1 complex). In **g,** the flipped W258 rotamer induced by PPSC-002 is shown as ball-and-sticks sitting in the PU.1 L73 pocket.

To characterize the binding mode of PPSC-001 and PPSC-002 with BAF60A and to understand the mechanism by which these compounds disrupt the interaction with PU.1, we co-crystallized each compound with BAF60A and determined the structures at 2.31 Å and 2.21 Å resolution, respectively. Clear electron density was observed for each compound in the hydrophobic core of BAF60A^YLD^ encompassing F270 and W258 with main-chain interactions flanking each pose for stabilization (Fig. 6d, e). Surprisingly, PPSC-002 sits slightly lower than PPSC-001, interacting with the loop connecting β4-β5, reaching both the amide NH of H259 and carbonyl amide of Q265, and occupies a unique cryptic pocket induced by the flipping of W258. This new conformational state adopted by W258, which has not been observed in prior structures, opens the hydrophobic core to help accommodate the triple ring system of PPSC-002. While a certain degree of overlap exists between the 2 compound binding sites, each compound occupies a distinct pocket within the hydrophobic core of BAF60A.

Consistent with our FP data showing that PPSC-001 inhibits the interaction of BAF60A and PU.1 with enhanced activity compared to PPSC-002, superimposition of these compound structures with our PU.1-BAF60A complex structure shows a larger extent of overlap between the PPSC-001 binding site and PU.1 compared with PPSC-002 (Fig. 6f, g). Specifically, PPSC-001 overlays with 5 residues of PU.1 (T71-S75), with the phenyl bromide moiety protruding into the hydrophobic space occupied by L73 side chain of PU.1, and the amide overlaying with its peptide backbone. PPSC-002 only overlaps with residue T71, but the additional W258 flip induced by the compound also would sterically clash with L73. Since L73 is critical for BAF60A binding to PU.1, this minimal overlap is sufficient to disrupt the interaction. Overall, both compounds occupy one of the critical interaction hot spots at L73 of the 70-78 loop of PU.1 and exhibit 2 unique binding modes capable of disrupting the PU.1-BAF60A interaction.

## Discussion

The BAF chromatin remodeling complex is a critical effector of TF regulation, and while many studies have established the close interplay between TFs and BAF^25^, there is a dearth of evidence and understanding of the direct, physical interaction between TFs and BAF. Here we establish, for the first time, the direct interaction of the human pioneer TF PU.1 with the BAF complex. We begin with a comprehensive set of cellular and functional genomic work to establish how PU.1 and BAF functionally cooperate for the regulation of transcription in AML cell lines. Next, through a series of biochemical and biophysical approaches, we measure and map the physical interaction of PU.1 and cBAF and provide a structural understanding of how PU.1 interacts with the YLD+SWIB of BAF60A via a high-resolution crystal structure.

YEATS domains bind acetylated (Ac) or crotonylated (Cr) histone tails^60^, and have been identified as therapeutic targets. As an example, the YEATS domain-containing protein ENL is a therapeutic target in MLL-fusion-positive AML. Because BAF60 contains a YEATS-like domain, it may have been expected to also be a histone reader domain. However, our data demonstrates that the YLD+SWIB domain in BAF60 has been repurposed as a protein-protein interaction domain, engaging with a disordered stretch within PU.1 but not recognizing a modified histone peptide. Similar to the functional role played by the ENL-histone-Ac interaction in AML, our work highlights the physiological importance of the interaction between BAF60A^YLD+SWIB^ and PU.1^70–100^ in AML. Additionally, our data suggests that PU.1 may be able to engage all 3 classes of BAF complex (ncBAF, cBAF, PBAF) through their shared BAF60 subunit. Future studies may reveal how each class works with PU.1 to regulate transcription in hematologic lineages.

Despite the many approaches to drug TFs, to date, the ones that have been targeted directly, such as nuclear hormone receptors (e.g. androgen^71^ and estrogen receptors), HIF1llJ^76^, and TEAD, possess well-defined druggable pockets. Notably, the HIF1llJ and TEAD inhibitors are PPI disruptors, preventing the interaction with key binding partners for therapeutic benefit. However, most TFs are still considered “un-druggable” due to the absence of well-defined, druggable pockets, given their high degree of intrinsic disorder^79^. Here we present a strategy for drugging the PPI between TFs and chromatin remodelers by leveraging the ligandable sites on BAF. Our HTS screen identifies compounds that bind to BAF60, disrupt the PU.1-BAF60 interaction in vitro, and provide a potential starting point for therapeutic development in AML.

Because TF–BAF interactions can be tissue-specific, driven by the expression patterns of both the TF and the BAF subunit paralogs, this strategy would allow modulation of TF activity in a context-dependent manner, and provide a high degree of selectivity for the TF of interest. In our example, the restricted expression of PU.1 makes the interaction of PU.1 with BAF60A critical in hematopoietic lineages and should allow precise targeting of AML cells in vivo. Furthermore, given the lack of sequence conservation in IDRs, even within members of the ETS family of TFs, disrupting a specific TF–BAF interaction is likely to yield a high degree of target selectivity. Overall, sequence divergence in IDRs, coupled with their structural plasticity, means that each unique TF–BAF interface could be targeted by a corresponding PPI inhibitor, potentially allowing development of a multitude of cell- or tissue-specific therapies for TF-driven pathologies, each with a substantially reduced risk of off-target effects.

Together, our work establishes a modular and scalable platform for drugging transcription factors through disruption of their interactions with chromatin remodelers. By integrating cell-based functional genomics, biochemical and biophysical studies, structural biology, and high-throughput screening, we have developed a comprehensive workflow for identifying, validating, and targeting TF–BAF interactions. Future applications of this workflow across TF families and chromatin remodeler subunits could unlock a new class of targeted therapies for transcriptionally addicted cancers and beyond.

## Methods

### PRISM Screen

A pooled cell viability assay was performed using PRISM (Profiling Relative Inhibition Simultaneously in Mixtures) as previously described^39^ with the following modifications. Cell lines were obtained from the Cancer Cell Line Encyclopedia (CCLE) collection and adapted to RPMI-1640 medium without phenol red, supplemented with 10% heat-inactivated fetal bovine serum (FBS), in order to apply a unique infection and pooling protocol to this large compendium of cell lines. A lentiviral spin-infection protocol was executed to introduce a 24-nucleotide barcode in each cell line, with an estimated multiplicity of infection (MOI) of 1 for all cell lines, using blasticidin as selection marker. Over 750 stably barcoded PRISM cancer cell lines were then pooled by doubling time, in pools of 25. For the screen execution, instead of plating a pool of 25 cell lines^39^, all the adherent or all the suspension cell line pools were plated together using T25 flasks (100,000 cells/flask) or 6-well plates (50,000 cells/well), respectively. Cells were treated with either DMSO or FHT-1015 in an 8-point 3-fold dose-response concentration range in triplicate, starting from a top concentration of 10 µM (14 nM to 10 µM). As a positive control, cells were treated in parallel with 2 previously validated compounds known to reduce cell viability, the pan-Raf inhibitor AZ-628 and the proteasome inhibitor bortezomib, using a top concentration of 2.5 mM and 0.039 mM, respectively. After 3 days of exposure, cells were lysed, genomic DNA was extracted, and barcodes were amplified by PCR and detected with next-generation sequencing. Cell viability was determined by comparing the counts of cell line specific barcodes in FHT-1015- and positive control-treated samples to those at day 0 and those treated with DMSO. Dose-response curves were fit for each cell line and corresponding area under the curve (AUC) values were calculated. AUC values were displayed as box plots and cell lines with AUCs less than the median were considered sensitive. For each box, edges of the box represent 25^th^ and 75^th^ percentiles; vertical lines represent median values; horizontal lines represent minimum and maximum values.

### ChIP-seq

Chromatin immunoprecipitation sequencing (ChIP-seq) assays were carried out on THP-1 cells treated with DMSO or 100 nM Compound 49 for 24h. Cells were cross-linked for 15 min in 1% formaldehyde at room temperature and quenched in 125 mM glycine for 5 min. Fixed cells were shipped to Active Motif where ChIP-seq was performed. In brief, cells were fixed with 1% formaldehyde for 15 min and quenched with 0.125 M glycine. Chromatin was isolated by adding lysis buffer, then disrupted with a Dounce homogenizer. Lysates were sonicated and the DNA sheared to an average length of 300 to 500 bp with the Active Motif EpiShear probe sonicator (part #53051) and cooled sonication platform (53080). Genomic DNA (Input) was prepared by treating aliquots of chromatin with RNase, proteinase K, and heat for de-crosslinking, followed by ethanol precipitation. Pellets were resuspended and the resulting DNA was quantified on a NanoDrop spectrophotometer. Extrapolation to the original chromatin volume allowed quantitation of the total chromatin yield. An aliquot of chromatin (30 µg) was precleared with protein A agarose beads (Invitrogen). Genomic DNA regions of interest were isolated using 10 µL of antibody against BRG1 (Abcam, AB110641), 20 µL of antibody against PU.1 (Santa Cruz, sc-352), or 4 µL of antibody against H3K27Ac (Active Motif, 39133). Complexes were washed, eluted from the beads with SDS buffer, and subjected to RNase and proteinase K treatment. Crosslinks were reversed by incubating overnight at 65^°^C, and ChIP DNA was purified by phenol-chloroform extraction and ethanol precipitation. Illumina sequencing libraries were prepared from the ChIP and Input DNAs using the standard consecutive enzymatic steps of end-polishing, dA-addition, and adaptor ligation using the Active Motif custom liquid handling robotics pipeline. After the final 15 cycle PCR amplification step, the resulting DNA libraries were quantified and sequenced on an Illumina NexSeq 500. Single-end sequencing was performed with read length of 75 bp at a depth of 30M reads on the Illumina platform. Reads were aligned to the human genome (hg19) using BWA mem with the default parameters. Alignments with a quality lower than 10 were discarded using samtools and duplicated alignments were removed with the picard MarkDuplicates tool. Peaks were called with the Macs2 callpeak function and fold enrichment coverage tracks were created using the bdgcmp function of the same package. Peaks that were on the ENCODE blacklist of known false ChIP-Seq peaks were removed.

### ATAC-seq

Cells were treated with Compound 49 at 100 nM or DMSO for 24h. ATAC-seq was then performed in biological triplicate as previously described. After library generation, left- and right-size selection was performed with SPRIselect beads (Beckman Coulter) according to the manufacturer’s recommendations. Sequencing was performed using the Illumina HiSeq3000 platform and approximately 60M of 150 bp paired-end reads were obtained per sample. Sequence analysis was performed as follows. Reads were trimmed for adapter sequences using cutdapt and aligned to version hg38 of the human genome using bowtie2 (parameters -k 4, -X2000, --local,--mm). Multimapping alignments were removed and only the alignment with the highest score for each read pair was kept. Duplicate reads were removed using Picard Tookit and alignments mapping to the mitochondrial genome were excluded. Alignments were filtered for size smaller than 120 bp and read ends shifted +4 or −5 nucleotides depending on the direction of the read. Peak calling was performed using MACS2.0 with a shift parameter of −75 and an extension size of 150 without modelling. Signal tracks were created using the fold enrichment option. Consistent peaks among replicates were obtained using irreproducible discovery rate (IDR) with a soft IDR threshold of 0.1. The consensus peak list considered was the one containing the largest number of regions after pairwise IDR analysis of all replicates. Differential genome accessibility calculation was performed using limma package. Motif enrichment was performed using Meme suite tools (MEME-chip) and CISBP transcription motif data from the MEME motif database, version 12.19. Ridge regression analysis was performed using glmnet R package with coefficients calculated using the minimum estimated lambda. Genomic context annotation was performed using the annotatePeaks function of HOMER package using the hg38 version of the human genome.

### RNA-seq

MV-4-11 cells were treated with or without 100 nM FHT-1015 for 24h, or with doxycycline treatment for 48h for shRNA experiments and THP-1 cells were treated with 100 nM Compound 49 for 24h. Total RNA was extracted from frozen cell pellets and washed with phosphate-buffered saline (PBS). RNA (1 μg per sample) was used as input material. Sequencing libraries were generated using NEBNext Ultra RNA Library Prep Kit for Illumina (NEB, USA) following the manufacturer’s recommendations, and index codes were added to each sample. mRNA was purified from total RNA using poly-T oligo-attached magnetic beads. Fragmentation was carried out using divalent cations under elevated temperature in NEBNext First Strand Synthesis Reaction Buffer (5X). First strand cDNA was synthesized using random hexamer primer and M-MuLV Reverse Transcriptase (RNase H-). Second strand cDNA synthesis was performed using DNA Polymerase I and RNase H. Remaining overhangs were converted into blunt ends via exonuclease/polymerase activities. After adenylation of 3 ends of DNA fragments, NEBNext Adaptor with hairpin loop structure were ligated to prepare for hybridization. To select cDNA fragments of preferentially 150 to 200 bp in length, the library fragments were purified with AMPure XP system (Beckman Coulter). Then 3 μL of USER Enzyme (NEB) was used with size-selected, adaptor-ligated cDNA at 37°C for 15 min followed by 5 min at 95°C before PCR. PCR was performed with Phusion High-Fidelity DNA polymerase, Universal PCR primers, and Index (X) Primer. Finally, PCR products were purified (AMPure XP system) and library quality was assessed on the Agilent Bioanalyzer 2100 system. Sequencing reads were aligned to version hg38 of the human genome using gencode v30 gene annotations. STAR aligner version 2.0.2 with the following parameters was used: twopassMode Basic, outFilterMultimapNmax 20, alignSJoverhangMin 8, alignSJDBoverhangMin 1, outFilterMismatchNmax 999, outFilterMismatchNoverLmax 0.1, alignIntronMin 20, alignIntronMax 1000000, alignMatesGapMax 1000000, outFilterType BySJout, outFilterScoreMinOverLread 0.33, outFilterMatchNminOverLread 0.33, limitSjdbInsertNsj 1200000, outSAMstrandField intronMotif, outFilterIntronMotifs None, alignSoftClipAtReferenceEnds Yes, quantMode TranscriptomeSAM GeneCounts, outSAMtype BAM Unsorted, outSAMunmapped Within, chimSegmentMin 15, chimJunctionOverhangMin 15, chimOutType Junctions WithinBAM SoftClip, chimMainSegmentMultNmax 1, outSAMattributes NH HI AS nM NM choutSAMattrRGline ID:rg1 SM:sm1. Differential gene expressions were calculated using the limma-voom R package. For shPU.1 differential gene expression analysis, non targeting control (NTC) without doxycycline was used as the negative control. All differentially expressed genes are statistically significant (padj < 0.05) and are at least 2-fold downregulated or upregulated, except no logFC cutoff was applied in shPU.1 differential gene expression analysis. Gene set enrichment analysis and gene ontology term overrepresentation analysis were performed using clusterProfiler and DOSE R packages.

### Purification of PU.1 and mutants

The full-length open reading frame (ORF) of PU.1 was subcloned into pcDNA3.4 vectors with a C-terminal MBP (maltose binding protein) tag separated by a TEV (tobacco etch virus) cleavage sequence. MBP-tagged full-length PU.1 was used for all experiments unless a truncation was specified. The plasmid was mixed with polyethyleneimine (PEI) at a 1:6 ratio in Opti-MEM medium (Thermo Fisher Scientific) and transfected into HEK293-F cells at a final DNA concentration of 1 mg/L. Cultures were incubated at 37°C, 130 rpm, 8% CO_2_ for 72h. Cells were harvested by centrifugation and the cell pellet was resuspended in lysis buffer (25 mM Tris, pH 8.0, 500 mM NaCl, 1 mM DTT, 5% glycerol, 10 mM MgCl_2_, 1 μg/mL DNase, Protease Inhibitor Cocktail III [Thermo Fisher Scientific], 0.5% Triton X-100). Cells were lysed by sonication (3 sec on, 5 sec off, 10 min) and lysate was clarified by centrifugation (30,000 rpm, 30 min, 4°C). Supernatant was passed over an amylose column (New England Biolabs) and washed with binding buffer (25 mM Tris, pH 8.0, 500 mM NaCl, 1 mM DTT, 5% glycerol). Immobilized protein was eluted off the column with binding buffer supplemented with 20 mM maltose. Eluted protein was loaded on a Q-HP column (Cytiva) and washed with binding buffer (25 mM Tris, pH 7.5, 100 mM NaCl, 1 mM DTT, 5% glycerol). Bound protein was eluted with a 100 to 500 mM NaCl gradient over 24 column volumes (CV). Eluted protein was further purified via SEC on a Superdex 200 increase column (Cytiva) in 25 mM Tris, pH 8.0, 300 mM NaCl, 1 mM DTT, 5% glycerol. Peak fractions were collected and concentrated in a 30 kDa cutoff concentrator (Amicon Ultra, Millipore) and analyzed for purity by SDS-PAGE. PU.1 mutant constructs were purified similarly.

### Purification of BAF complex and subcomplex

Endogenous BAF complex was purified as described previously^48^. For recombinantly purified BAF, ORFs for human BAF complex subunits (BRG1, BRM, ARID1A/B, BAF60, BAF170/155, BAF47, BAF57, BAF45D, ACTB, ACTL6A, BCL7A, and SS18) were subcloned into pcDNA3.4 vectors with no tag or a FLAG-tag followed by a 3C protease cleavage sequence. Full BAF consisted of plasmids encoding BRG1, ARID1A (991-2285), BAF170, BAF155, BAF47, BAF60A, BAF57, BAF45D, ACTB, ACTL6A, BCL7A, and SS18.

The structural core consisted of plasmids encoding for BRG1 N-terminus (1-465), ARID1B (1028-2236), BAF60A, BAF170, BAF47, BAF57, and BAF45D. The catalytic core contained BRM (416-1326) and ARP module subunits, BAF53A, ACTB, and BCL7A. The plasmids were mixed with PEI at a 1:6 ratio in Opti-MEM medium (Thermo Fisher Scientific) and transfected to Expi293F cells at a final DNA concentration of 3 mg/L. Cultures were incubated at 37°C, 125 rpm, 8% CO_2_ for 72h. Cells were harvested by centrifugation and the cell pellet was resuspended in lysis buffer (50 mM HEPES, pH 8.0, 300 mM NaCl, 0.25% CHAPS, 2 mM MgCl2, 0.25 mM EDTA, 2 mM DTT, 1 mM PMSF, cOmplete protease inhibitor cocktail (Roche)). Cells were lysed by Dounce homogenizer (Type A, 40 strokes) on ice and lysate was clarified by ultracentrifugation (17,000 rpm, 30 min, 4°C). Lysate was incubated with Anti-FLAG M2 resin (Sigma) for 3h at 4°C. Resin was transferred to gravity column, washed with lysis buffer, and bound protein was eluted with 50 mM HEPES, pH 8.0, 300 mM NaCl, 0.25% CHAPS, 2 mM MgCl2, 0.25 mM EDTA, 2 mM DTT, 250 µg/ml FLAG peptide. Eluted protein was loaded onto MonoQ column (Cytiva) and washed with 20 mM HEPES, pH 7.5, 100 mM NaCl, 10% Glycerol, 0.5 mM TCEP. Protein was eluted with a 100-500 mM NaCl gradient over 15 CV. Eluted protein was further purified via SEC on a Superose 6 increase column (Cytiva) in 20 mM HEPES, pH 7.5, 200 mM NaCl, 2% Glycerol, 1 mM TCEP. Peak fractions were collected and concentrated in 100 kDa cutoff concentrator (Amicon ultra, Millipore) and analyzed for purity by SDS-PAGE.

### Purification of BAF60A and mutants

BAF60A constructs and truncations were expressed from plasmid pET21b-His-FLAG-GSGS-TEV-S-SMARCD1 in *E. coli* BL21(DE3) gold strain using IPTG induction. Briefly, cells were grown in 2 L LB media with addition of a 20 mL starter culture and 100 μg/mL ampicillin and incubated on an orbital shaker for 4-5h at 37°C. Once cells reached OD600 of ∼0.6, IPTG was supplemented at 0.1 mM to the cultures and temperature was reduced to 16°C for 16h before cells were harvested. Pellets from 10 L cultures were resuspended by addition of lysis buffer (∼10 mL/g cells) containing 20 mM HEPES, pH 7.5, 300 mM NaCl, 10% Glycerol, 1 mM TCEP. TCEP was not included in SWIB purification. The resuspension was sonicated on ice at 400W, 3 sec on, 3 sec off, for 25 min before clarified lysate was obtained by centrifugation. The supernatant was passed on a 15 mL Ni^2+^ affinity column preequilibrated with lysis buffer containing Ni-NTA beads from Qiagen. A series of wash steps were performed with lysis buffer and lysis buffer supplemented with 20 mM imidazole until a stable protein signal was reached. BAF60A protein was then eluted with lysis buffer containing 250 mM imidazole. The fractions containing BAF60A as determined by 15% SDS-PAGE gel were pooled and digested with his-TEV at a (10:1 ratio) overnight at 4°C dialyzed against lysis buffer. A second Ni^2+^ affinity column was performed with 3 mL column the next day collecting cleaved BAF60A in the flow-through and additional lysis buffer wash. Pooled protein was concentrated using Amicon Ultra-15 Centrifugal Filter Unit (10 kDa MW cutoff) before passing the sample through a final size exclusion column (Cytiva Superdex 75 16/600 GL) with final storage buffer containing 20 mM HEPES, pH 7.5, 300 mM NaCl, 5% Glycerol, 1mM TCEP.

For BAF60A^SWIB^, an additional cation exchange column (Cytiva HiTrap SP HP) was run before size exclusion. Protein was diluted into lysis buffer without salt and loaded onto the column preequilibrated with the same buffer A (20 mM HEPES, pH 7.0, 10% glycerol) at a flow rate of 3 mL/min. The column was washed with 5 CV of buffer A and protein was eluted using a linear gradient of 0-30% buffer B (20 mM HEPES, pH 7.0, 1 M NaCl, 10% glycerol) for 10 CV. Fractions spanning a peak at 17% buffer B were collected and concentrated for size exclusion. Final buffer contained 20 mM HEPES, pH 7.0, 300 mM NaCl, 5% glycerol.

### Purification of ENL^YEATS^

The ORF corresponding to human ENL^YEATS^ residues 1-148 was subcloned into the pET21b vector with a C-terminal 6x His-tag and expressed in *E. coli* BL21(DE3) gold strain. 10 L cultures were grown at 37°C to OD₆₀₀ ≈ 0.6, induced with 0.1 mM IPTG, and shifted to 16°C for 16h. Cultures were harvested by centrifugation, and the cell pellet was resuspended in lysis buffer (50 mM Tris pH 7.5, 500 mM NaCl, 20 mM imidazole, 2 mM DTT). The resuspension was sonicated on ice at 400W, 3 sec on, 3 sec off, for 25 min and lysate was clarified by centrifugation. Supernatant was loaded onto a 5 mL Ni^2+^ affinity column (Qiagen), washed with lysis buffer, and eluted off the column with a 20-300 mM imidazole gradient over 6 CV Fractions containing ENL^YEATS^ were pooled and further purified via SEC on a Superdex 75 increase column (Cytiva) in 25mM Tris pH 7.5, 500 mM NaCl, 2 mM DTT. Peak fractions were collected and concentrated in 10 kDa cutoff concentrator (Amicon ultra, Millipore) and analyzed for purity by SDS-PAGE.

### Crystallization of BAF60A apo, BAF60A:PU.1 complex, and BAF60A:compounds

BAF60A apo crystals were grown in sitting drop format using premade solution consisting of 20% w/v PEG 3350, 0.2 M NH_4_NO_3_ from the JCSG+ suite (Molecular Dimensions). Protein drops were prepared by mixing 28.8 mg/mL protein and crystallization buffer at a 2:1 ratio and incubating for 1 day at 18°C.

BAF60A:PU.1 complex was formed by incubating 10 mg/mL BAF60A^YLD+SWIB^ with PU.1^60–100^ at a 1:1 molar ratio for 2h at 4°C before co-crystallization. For PU.1^60–100^, the protein was expressed as with other PU.1 constructs and purified via the same steps as BAF60A described above including the TEV protease cleavage step. The final protein was concentrated with an Amicon Ultra-15 Centrifugal Filter Unit (3 kDa MW cutoff) for crystallization. The complex was screened against a series of ten sparse-matrix crystallization conditions in sitting drop mode (0.2 µL:0.2 µL complex-to-well solution) using NT8 crystallization robot from Formulatrix at 18°C. Microcrystalline clusters were observed in condition containing 0.1M HEPES sodium salt pH 7.5, 2.0 M ammonium sulfate, 2% v/v PEG MME 550 from JBScreen Kinase HTS (Jena Bioscience) and further optimized by varying pH and buffer system and subsequently supplementing the condition with additives and seeds. The drop from the final condition was set-up with 0.4 µL complex at 8.5 mg/mL (1:1 molar ratio), 1.0 µL crystallization buffer (0.1 M tri-sodium citrate pH 6.48, 1.97 M ammonium sulfate, 2% v/v PEG MME 550, 0.01 M taurine), 0.14 µL additive (10 mM EDTA final concentration), and 50 nL seeds. Drops were equilibrated against 40 µL of crystallization buffer at 18°C in SWISSCI MRC 3 sitting drop trays and crystals were observed after 3 days.

BAF60A:compound co-crystals were screened for initial crystallization conditions by incubating 20 mg/mL BAF60A(144-397) with compound at a 1:10 protein-to-compound molar ratio at 4°C for 2h and setting up 5 sparse-matrix crystallization screens using microseed matrix screening with apo seeds at 18°C (0.2 µL:0.2 µL:50 nL complex-to-well solution: seeds). BAF60A(144-397) apo seeds were prepared by setting up protein at 25 mg/mL via hanging drop at a 1:1 well ratio in crystallization buffer containing 0.1 M NaCl, 0.1 M Tris pH 8.5, 13% w/v PEG 10,000, 20% v/v glycerol at 18°C. Briefly, one crystal was diluted into 2 µL buffer and grinded into microseeds before diluting 1000-fold for screening. After 13 days, crystals were observed and directly harvested from screening plate in conditions 0.2 M ammonium formate pH 6.6, 20% w/v PEG 3350 for PPSC-001 and 0.2 M potassium fluoride pH 7.3, 20% w/v PEG 3350 for PPSC-002, which were both found in the PEG/ion screen from Hampton Research.

Complexed and apo crystals were cryo-protected with crystallization buffer supplemented with 15% glycerol (ethylene glycol was used for BAF60A:PU.1^60–100^ complex) before they were harvested and cryo-plunged in liquid nitrogen for data collection. BAF60A apo structure was determined using MrBUMP automated molecular replacement in CCP4. BAF60A:PU.1^60–100^ and compound structures were determined by molecular replacement in Phaser using either the Alphafold2 model of BAF60A or apo structure, as a search model. Subsequent rounds of model building and refinement were done using Coot and CCP4. Final data collection and refinement statistics are reported in Table 1. Structural figures were prepared using PyMOL (Schrödinger, LLC) with refined crystallographic models.

### SEC-MALS

Size exclusion chromatography with multi-angle light scattering (SEC-MALS) was used to determine the molecular weight of purified recombinant cBAF, cBAF structural core and cBAF catalytic core. A Superose 6 Increase 10/300 GL column (Cytiva) was equilibrated with 20 mM HEPES pH 7.5, 200 mM NaCl, 1 mM TCEP at 0.5 mL/min. 100 µg of protein was injected onto the column and eluent was passed through a Wyatt DAWN TREOS multi-angle light scattering detector and an Optilab T-rEX refractive index detector (Wyatt Technology). The data were analyzed using ASTRA software (Wyatt Technology) and the molecular weight was calculated using a dn/dc value of 0.185 mL/g.

### Biochemical pull-down assay

Amylose resin (New England Biolabs) was washed with buffer A (20 mM HEPES, pH 7.4, 100 mM NaCl, 0.4 mM TCEP) and 1 mg of MBP-tagged PU.1 constructs were immobilized per 1 mL of resin. PU.1 constructs were incubated for 15 min, 20°C, rotating in Eppendorf tubes. Resin was transferred to Micro-Spin columns (Thermo Fisher Scientific) and spun at 2000 x g for 30 sec. The resin was washed with 10 CV of buffer A. cBAF, cBAF structural core, cBAF catalytic core or BAF60A^YLD+SWIB^ were dialyzed and diluted in buffer A to a final concentration of 10 µM and added to the column. After incubation (30 min, 20°C), unbound protein was washed with 15 CV of buffer A. Bound protein was eluted with buffer A supplemented with 10 mM maltose. Eluate was analyzed by 4-12% SDS-PAGE gel.

### Analytical ultracentrifugation (AUC)

Sedimentation velocity analytical ultracentrifugation (AUC) experiments were performed using a Beckman XL-I centrifuge. Samples were sedimented in 2-channel sector shaped cells with sapphire windows and sedimentation was monitored using absorbance optics at 280 nm. Sedimentation was performed in buffer (20 mM HEPES, 150 mM NaCl, 0.05% Tween-20, 1 mM TCEP, pH 7.5). Zeba desalting columns were used immediately prior to loading the AUC cells to buffer match samples to reference buffer. Samples were loaded into the centrifuge and allowed to equilibrate to 25°C for 1h prior to the sedimentation of the samples. Samples were spun at 40,000 rpm (PU.1 alone) or 25,000 rpm (cBAF or cBAF+PU.1) and absorbance scans were collected continuously. cBAF concentration was held constant at 0.1 µM and PU.1 was titrated from 0.2 to 4 µM.

Absorbance data was fit using Sedfit to obtain continuous c(s) distributions. Weight average sedimentation coefficients of PU.1:cBAF were extracted and plotted against PU.1 concentration using GraphPad Prism (version 10.0.2) and fit to 1-site binding equation to obtain equilibrium binding parameters

### Mass spectrometry-based protein footprinting and analysis

Both PU.1 and cBAF were prepared in 1×PBS, pH 7.4, buffer. To prepare the cBAF-PU.1 complex, cBAF and PU.1 proteins were mixed at a 1:5 mol:mol ratio. The protein concentrations of cBAF and PU.1 were adjusted to 0.5 and 2.5 μM, respectively. Protein samples were mixed with EDC (240 mM) and GEE (8 mM) in PBS for 0, 2.5, 5, and 7 min at room temperature and quenched with 8 M urea and 200 mM DTT in 1 M ammonium acetate. Samples were cleaned, digested, and analyzed as described in Martinez et al ^48^.

For CF3-labeling, free cBAF and cBAF-PU.1 complex were exposed to x-ray in the presence of 30-mM sodium triflinate for 25, 40, and 55 msec and aluminum attenuation of 76-µm. All samples were cleaned, digested, and analyzed similarly to the EDC/GEE protocol.

### Isothermal titration calorimetry (ITC)

Proteins were dialyzed using Slide-A-Lyzer MINI Dialysis Devices, 10K molecular weight cutoff (MWCO) (Millipore Sigma, part #69570) overnight at 4°C in ITC buffer (20 mM HEPES pH 7.4, 100 mM NaCl, 0.4 mM TCEP, 5% glycerol). The dialyzed protein was centrifuged at 13,000 rpm at 4°C for 10 min. The supernatant was collected, and protein concentration was measured using a Thermo Scientific NanoDrop One Microvolume UV-Vis Spectrophotometer. The proteins were diluted to the desired concentrations (approximately 40 µM PU.1 constructs and 400 µM BAF60A constructs) in ITC buffer and loaded into a PEAQ-ITC machine (Malvern Panalytical). BAF60A^YLD+SWIB^ was titrated from syringe into the reaction cell containing PU.1 constructs at 20°C, stirring at 500 rpm. Experiments were performed using reference power at 10 µcal/sec, feedback at high, and a 60 sec initial delay. Data was analyzed using MicroCal PEAQ-ITC Analysis Software (version 1.41) with a 1-site model.

### Labeling of PU.1 peptide (PU.1^60–100^^TMR^) for FP assay

BODIPY TMR C5 Maleimide (Invitrogen BODIPY TMR C_5_-Maleimide, part #B30466) was solubilized at 10 mM in DMSO. PU.1 (60C-100) peptide was purified as was done for other PU.1 constructs, with an additional cleavage step wherein the MBP tag was cleaved with TEV protease for 1.5h at room temperature and flown through a Ni-FF column (Cytiva). The peptide was further purified with a Superdex 75 column (Cytiva) and analyzed for purity by SDS-PAGE. The peptide was labeled in labeling buffer (20 mM HEPES pH 7.0, 0.2 mM TCEP, and 100 mM NaCl) at a 10:1 molar ratio of dye to PU.1 for 2h at room temperature in the dark. The labeling reaction was quenched with DTT at a final concentration of 1 mM. Excess dye was removed using a Millipore Amicon Pro Purification System with a 3 kDa MWCO Ultra-0.5 device following the manufacturer’s protocol. Samples were exchanged into desalting buffer (20 mM HEPES pH 7.0, 100 mM NaCl, 1 mM DTT, 2% glycerol, and 0.001% Tween20). Labelled peptide was stored at −80°C.

### Fluorescence polarization assay

For FP binding experiments, 20 nM PU.1^60–100^^TMR^ peptide was incubated for 30 min with increasing concentrations of specified BAF60A constructs in triplicate. Assay was performed in black 384-shallow well Proxiplate Plus plates with a 9 µL total assay volume in assay buffer (20 mM HEPES pH 7.5, 100 mM NaCl, 0.5 mM TCEP, 0.005% Tween20, and 0.005% BSA) at room temperature in darkness. Microplates were spun for 1 min at 1,000 rpm and read on a PHERAstar FSX plate reader (BMG Labtech) using the TMR FP optic module. The resulting mP values were plotted against the protein concentration and fitted to a 4-parameter dose-response curve using GraphPad Prism (version 10.0.2) to obtain equilibrium dissociation constants (K_D_).

For FP competition experiments, 400 nM BAF60A^YLD+SWIB^ was incubated for 30 min with increasing concentrations of specified PU.1 constructs or compounds in triplicate. PU.1^60–100^^TMR^ peptide (20 nM) was added to the plates and incubated for an additional 30 min in darkness. Microplates were spun for 1 min at 1,000 rpm and read on a PHERAstar FSX plate reader (BMG Labtech) using the TMR FP optic module. The resulting mP values were plotted against the protein concentration and fitted to a 4-parameter dose-response curve using GraphPad Prism (version 10.0.2) to obtain half maximal inhibitory concentration (IC_50_) values. The assay was performed in black 384-shallow well Proxiplate Plus plates in 9 µL total assay volume in assay buffer (20 mM HEPES pH 7.5, 100 mM NaCl, 0.5 mM TCEP, 0.005% Tween20, and 0.005% BSA) at room temperature in darkness. All PU.1 constructs were buffer exchanged using 0.5 mL of 7 kDa Zeba desalting columns per manufacturer instructions. Protein concentration was quantified using a Thermo Scientific NanoDrop One Microvolume UV-Vis Spectrophotometer.

For FP HTS assays, 1 µL of 2.25 µM (3× final concentration) of BAF60A^YLD+SWIB^ was added to all wells, followed by 1 µL of assay buffer (20 mM HEPES pH 7.5, 100 mM NaCl, 0.5 mM TCEP, 0.005% Tween20, 0.005% BSA, 500 μM EDTA, 500 μM EGTA and 0.25% DMSO) and 22.5 nL of test compounds at a final concentration of 30 µM. Compounds were added using Echo. Microplates were spun for 1 min at 1,000 rpm. After a 30 min incubation, 1 µL of 60 nM (3× final concentration) PU.1^60–100^^TMR^ was added to wells. Microplates were spun for 1 min at 1,000 rpm before reading on a PHERAstar FSX plate reader (BMG Labtech) using the TMR FP optic module. The assay was performed in a 3 µL total assay volume in black 1536 Greiner plates (part #782076) at room temperature. Data were normalized to the signal from the neutral control (DMSO only) and inhibition control (no BAF60A^YLD+SWIB^) to obtain percent inhibition. Hits (inhibitors), defined as compounds that exhibit percent inhibition of greater than 3 times the standard deviation (3SD=18%), were serially diluted 3-fold in a 10-point dose-response titration. The resulting percent inhibition values were plotted against the protein concentration and fitted to a 4-parameter dose-response curve using GraphPad Prism (version 10.0.2) to obtain half maximal inhibitory concentration (IC_50_) values.

### Surface plasmon resonance (SPR)

Histone peptide and compounds were evaluated for their binding to BAF60A^YLD+SWIB^ and ENL^YEATS^ by SPR using the Biacore S200 system (Cytiva) at 25°C. Both BAF60A^YLD+SWIB^ and ENL^YEATS^ were immobilized on a Series S CM5 Sensor chip (Cytiva) by amine coupling. The surface was activated with a 1:1 mix of N-hydroxysuccinimide (0.1 M) and 1-ethyl-3-(3-dimethylaminopropyl) carbodiimide (0.4 M) for 420 sec at 10 μL/min. Protein solutions prepared in 10 mM HEPES, pH 7.0, were pulsed on the surface for 360 sec (BAF60A^YLD+SWIB^) and 180 sec (ENL^YEATS^) at 10 μL/min and then quenched by 420 sec of ethanolamine (1 M, pH 8.0). A final bound response of ∼2000 RUs (BAF60A^YLD+SWIB^) and ∼1000 RUs (ENL^YEATS^) was obtained. Two- or 3-fold, 8-point titrations of compounds and H3K27Cr (APRKQLATKAARK^Cr^SAPATGG) were flowed over for association and dissociation times of 60 sec and 100 sec, respectively, at 30 μL/min in multicycle mode in SPR buffer (20 mM HEPES pH 7.5, 100 mM NaCl, 1 mM TCEP, 0.02% Tween20, 1% glycerol, 1% DMSO).

Data were analyzed using Biacore S200 Evaluation software. Sensograms were solvent-corrected, followed by reference and blank subtraction.

H3K27Cr peptide was synthesized at Biopeptide Co., Inc.

### Cell culture

Cell lines were purchased from ATCC (MV-4-11, THP-1, KASUMI-6, HEK293-T, HEL 92.1.7) and DSMZ (EOL-1, OCI-AML-2, OCI-AML-3, NOMO-1). Cell lines were grown according to the ATCC or DSMZ recommendations and were tested routinely to ensure that they were negative for mycoplasma.

### Cell proliferation assays

3- and 7-day proliferation assays were run in technical triplicate at Chempartner with the following cell lines: MV-4-11, THP-1, KASUMI-6, HEL 92.1, EOL-1, OCI-AML-2, OCI-AML-3, and NOMO-1. Cells were incubated at 37°C with a dose titration of compounds in 96-well plates for 3 days. On day 3, cells were split, and a portion of cells were incubated with Cell-Titer Glo 2.0 (Promega) to measure relative growth on day 3 using an Envision plate reader (Perkin-Elmer). Remaining cells were re-plated back to the starting density in fresh media and fresh compound and incubated at 37°C for an additional 4 days, after which relative growth on day 7 was measured as on day 3. Growth relative to a DMSO control was measured, curves were fit using a 4-point non-linear regression model, and absolute IC_50_ values were measured in GraphPad Prism (version 10.0.2).

### Immunoprecipitation and Western blots

HEK-293-T cells were transfected with plasmids expressing V5-tagged PU.1 by plating cells at 80% confluence prior to transfection using polyethylenimine (PEI) in a 3:1 PEI:DNA ratio. Cells were harvested after 72h. Nuclear extracts and immunoprecipitation were performed by washing cells with cold PBS and resuspended in a hypotonic buffer containing 50 mM Tris (pH 7.5), 0.1% NP-40, 1 mM EDTA, and 1 mM MgCl_2_ supplemented with protease inhibitors. Lysates were pelleted at 5,000 rpm for 5 min at 4°C. Supernatants were discarded and nuclei were resuspended in a high-salt buffer containing 50 mM Tris pH 7.5, 300 mM NaCl, 1% NP-40, 1 mM EDTA, and 1 mM MgCl_2_ supplemented with protease inhibitors. Lysates were incubated on ice for 10 min with occasional vortexing. Lysate was pelleted at 21,000 g for 10 min at 4°C. Supernatants were quantified and 1,000 μg of protein was used for immunoprecipitation with 3 μg of an anti-V5 antibody overnight at 4°C. Protein-G Dynabeads were added for 2h and washed with immunoprecipitation buffer. Beads were eluted with loading LDS and loaded onto SDS-PAGE for analysis. Membranes were incubated with antibodies targeting V5, PU.1, BRG1 (*SMARCA4*), and BAF170 (*SMARCC2*) (Cell Signaling Technology). Likewise, knockdown of PU.1 was measured in MV-4-11 cells after treating with doxycycline for 48h. Nuclear extracts were obtained using the same protocol as above and 500 μg of protein was run for Western blot analysis. Following incubation, protein levels were measured on an Odyssey CLx Imaging System (LI-COR).

### PU.1 shRNA and rescue experiments

The following SMARTvector Inducible Lentiviral shRNA, targeting PU.1 (*SPI1*), and a non-targeting control were acquired from Horizon Discovery:

ShPU.1#1 (v31hspgg_4718277) **Mature Antisense:** CCCGTCTTGCCGTAGTTGC

ShPU.11#2 (v31hspgg_5355474) **Mature Antisense:** TTGTCCACCCACCAGATGC

ShPU.11#3 (v31hspgg_8494863) **Mature Antisense:** GGGTACTGGAGGCACATCC

ShNonTargeting (shNT) Cat#: VSC6580

These shRNAs were packaged into lentivirus and infected into MV-4-11 cells. After puromycin selection, cells were treated with doxycycline and PU.1 levels were assessed using Western blot. A V5-tagged PU.1-resistant ORF (PU.1r) with blasticidin resistance was created for rescue experiments as described below.

**Wild-type SPI1 (PU.1r) shRNA resistant sequence (shRNA target sequences underlined w/ changes capitalized):** atgttacaggcgtgcaaaatggaagggtttcccctcgtcccccctccatcagaagacctggtgccctatgacacggatctatac caacgccaaacgcacgagtattacccctatctcagcagtgatggggagagccatagcgaccattactgggacttccacccc caccacgtgcacagcgagttcgagagcttcgccgagaacaacttcacggagctccagagcgtgcagcccccgcagctgca gcagctctaccgccacatggagctggagcagatgcacgtcctcgatacccccatggtgccaccccatcccagtcttggccac caggtctcctacctgcccc**gCatgtgTTtGcaAtaTcc**atccctgtccccagcccagcccagctcagatgaggaggagg gcgagcggcagagccccccactggaggtgtctgacggcgaggcggatggcctggagcccgggcctgggctcctgcctgg ggagacaggcagcaagaagaagatccgcctgtaccagttcctgttggacctgctccgcagcggcgacatgaaggacT**CA atAtggtgggtTgaTaa**ggacaagggcaccttccagttctcgtccaagcacaaggaggcgctggcgcaccgctggggc atccagaagggcaaccgcaagaagatgacctaccagaagatggcgcgcgcgctgc**gAaaTtaTggAaaAacCgg** cgaggtcaagaaggtgaagaagaagctcacctaccagttcagcggcgaagtgctgggccgcgggggcctggccgagcg gcgccacccgccccactga

The PU.1r was then infected via lentivirus into shPU.1- or shNT-containing cells selected with blasticidin. Cell viability rescue and ORF expression were assessed using cell proliferation assays and Western blots.

### Test compounds

PPSC-001 and PPSC-002 were purchased from commercial sources and used without further purification.

Compound 94 was synthesized as in.

FHT-1015 and Compound 49 were synthesized as in^37^.

## Author contributions

A.M.T., S.F.B., G.J.S., D.S. conceptualized and designed the overall study. D.S., D.M., L.V.d.P., S.W., S.E.S., J.W.S., M.R.M., J.K., S.H., B.T. designed, executed and analyzed experiments. S.T., G.J.S., D.L.L. performed Bioinformatic Analysis. D.M., J.R.C., B.T., G.M.P., J.W.S, D.S., A.M.T., Y.W.D designed constructs and purified the proteins. J.R.C., G.M.P. determined the x-ray crystal structures and J.R.C., D.S., A.M.T, L.V.d.P., contributed to structural analysis and mutational studies. D.S., L.V.d.P., S.J.B., A.M.T., M.N. executed and analyzed High-Throughput Screening (HTS) data. W.F.A., D.H., Y.G., K.J.W. designed and synthesized compounds. D.S., J.R.C., S.T., G.J.S., A.M.T., Y.W.D. wrote the initial draft and A.K., J.L.P. reviewed and edited the final draft.

## Competing interests

The authors declare no competing interests.

**Correspondence and requests for materials** should be addressed to Asad M. Taherbhoy.

## Supplementary materials

Supplementary materials include 6 extended data figures and 2 extended data tables.

## Data availability

The coordinates and structure factor files for X-ray crystal structures reported in this manuscript have been deposited to the Protein Data Bank with identifiers 9PSJ (BAF60A^144–407^ apo), 9PSL (BAF60A^YLD+SWIB^ with PU.1 peptide), 9PSQ (BAF60A^144–397^ with PPSC-001), and 9PSR (BAF60A^144–397^ with PPSC-002). All sequencing data (RNA-seq, ATAC-seq, and ChIP-seq) generated in this study were deposited at Gene Expression Omnibus (GEO) under accession number GSE304789. Data used to generate Extended Data Fig. 1d were taken from GEO (accession number GSE241445).

## Supplementary Materials

### Extended data

**Extended Data Fig. 1.**
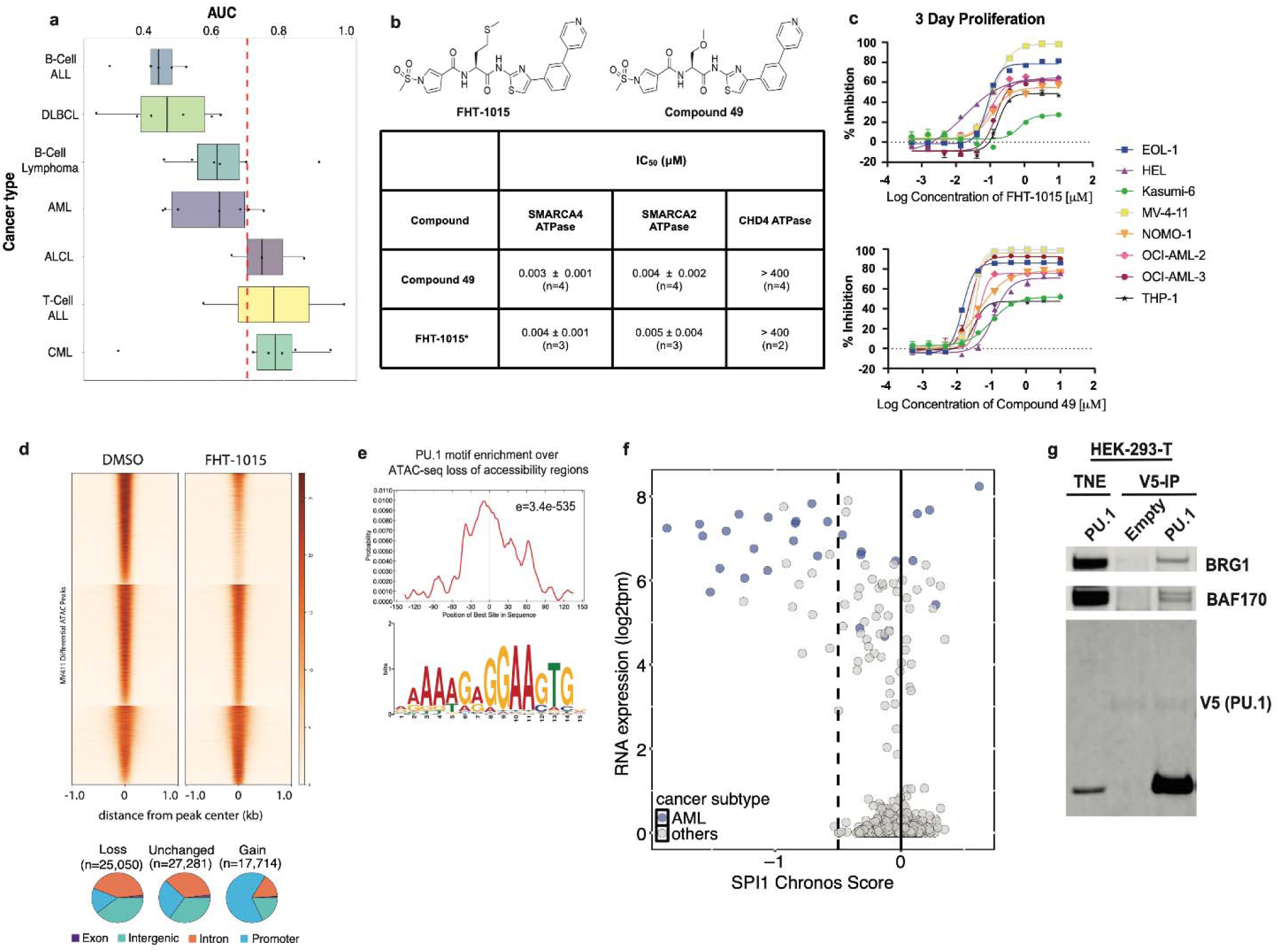
**a,** Box plot displaying AUC values of hematologic malignancy cell lines (n=35) treated with FHT-1015 for 3 days. Sensitivity threshold for FHT-1015 is indicated with the dashed red line. Values to the left of the sensitivity threshold indicate sensitive cell lines. **b,** Chemical structures of Compound 49 and FHT-1015 (*extracted from^34^). DNA-dependent ATPase activity was measured by the ADP-glo assay in the presence of inhibitor using full-length SMARCA4, SMARCA2, and CHD4. IC_50_ values (geometric mean ± SD) are shown. **c,** Percentage of proliferation inhibition in response to increasing doses of FHT-1015 (top graph) or Compound 49 (bottom graph) for 3 days in 8 leukemic cell lines (EOL-1, HEL, Kasumi-6, MV-4-11, NOMO-1, OCI-AML-2, OCI-AML-3, and THP-1). **d,** Tornado plot showing changes in chromatin accessibility (sorted by DMSO signal) in MV-4-11 cells in response to treatment with FHT-1015 for 4h. Pie charts showing distribution of genomic features of ATAC regions, by type of change in chromatin accessibility. Data for this analysis were obtained from GEO (GSE241445). **e,** Enrichment p-value and motif probability graph for SPI-1 as the most enriched motif at the loss-of-accessibility regions in MV-4-11. **f**, Relationship ￼*SPI1* (PU.1) Chronos score and *SPI1* RNA expression level across various cancer cell lines (n=1,108). AML cell lines are displayed in blue. Other cell lines are displayed in gray. Chronos score and RNA expression values were obtained from DepMap Public 25Q2 dataset (https://depmap.org/portal). Dashed line indicates the predicted sensitivity threshold of cell lines to the loss of *SPI1.* Cell lines to the left of the dashed line are considered sensitive based on DepMap Chronos Score. **g,** Nuclear protein input and anti-V5 immunoprecipitation (right) in HEK-293-T cells transfected with V5-PU.1 or empty vector.

**Extended Data Fig. 2.**
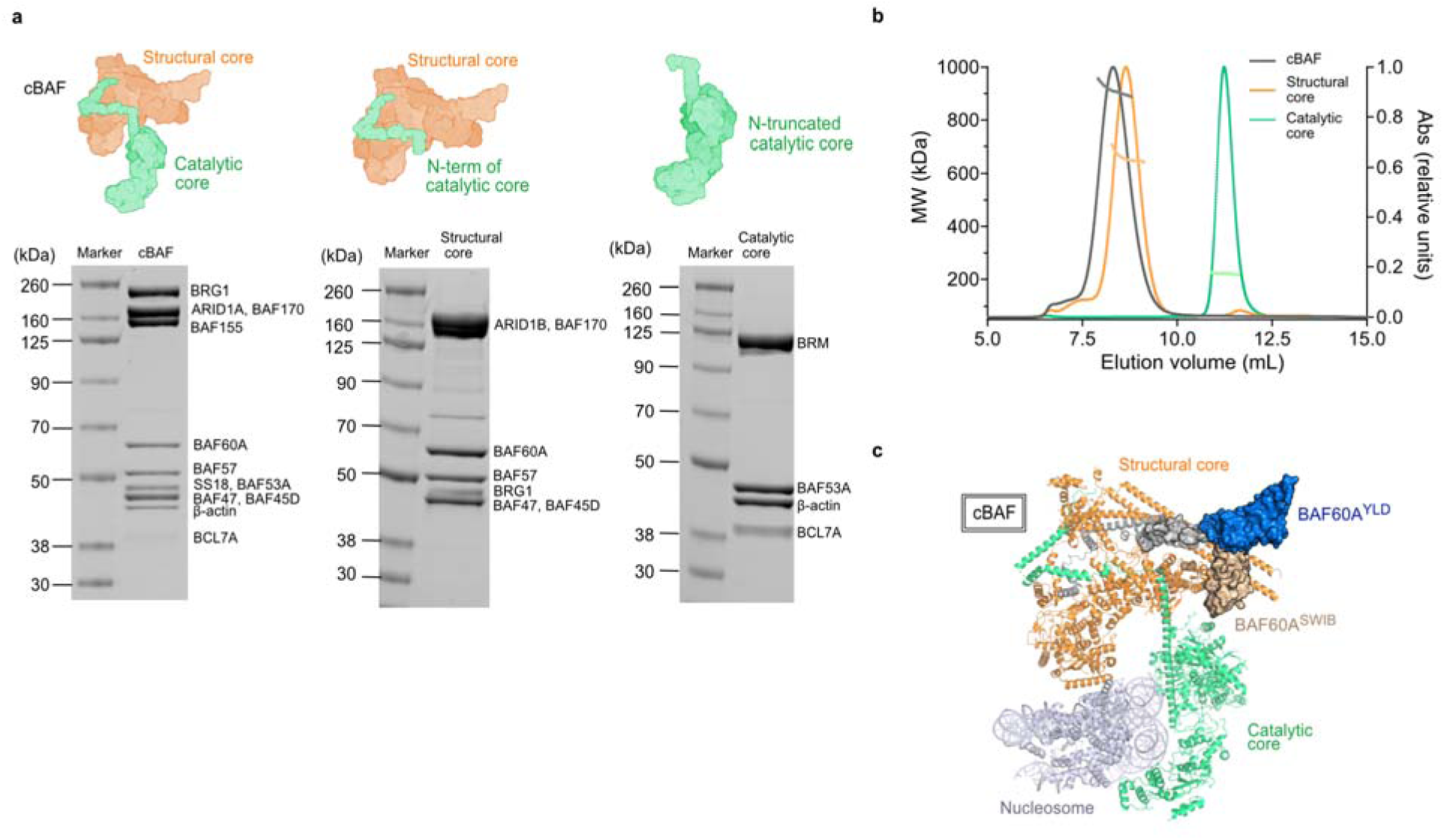
**a,** 4-12% SDS-PAGE gel of purified reconstituted full cBAF, structural core, and catalytic core with each component labeled. Densitometry indicated purity of >90%. **b,** SEC-MALS analysis of full cBAF (gray), structural core (orange), and catalytic core (green) of cBAF. SEC-MALS revealed a single, monodisperse peak corresponding to an absolute molecular weight of 911 kDa, 658 kDa, and 220 kDa for cBAF, its structural core, and its catalytic core, respectively, in agreement with the expected molecular weights of 938 kDa, 647 kDa, and 222 kDa. **c,** Superposition of BAF60A^YLD+SWIB^ apo structure with human cBAF complex (PDB code: 6LTJ)^56^ aligning by BAF60A^SWIB^. The structural core (orange), catalytic core (green), and nucleosome (gray) of cBAF and BAF60A^YLD^ (blue), BAF60A^SWIB^ (beige), and helical region of coiled coil (CC, dark gray) of BAF60A^YLD+SWIB^ apo are labeled. The resolved regions of BAF60A from cBAF are colored accordingly (BAF60A^SWIB^, beige; CC and pillar helix [pill], dark gray).

**Extended Data Table 1.** MS footprinting data with protection ratios for ARID1B peptides following EDC/GEE labeling of cBAF, suggesting interaction with PU.1. The peptides with the highest degree of protection are bolded.

| **Location** | **ARID1B sequence within cBAF** | **Site of modification** | **Protection ratio** |
| --- | --- | --- | --- |
| 1872-1889 | PGALPEDANPGPQTESSK | E1886, E1877, D1878 | 1.14 |
| **1905-1911** | **LLEDEPR** | **E1907, D1908, E1909** | **2.48** |
| 1997-2008 | DEWWWDCLEVLR | D2002, E2005 | 0.93 |
| **2076-2094** | **LSIQDNNVDLILATPPFSR** | **D2080, D2084** | **1.28** |
| 2200-2208 | SEFLLHEGR | E2201, E2206 | 0.76 |

**Extended Data Fig. 3.**
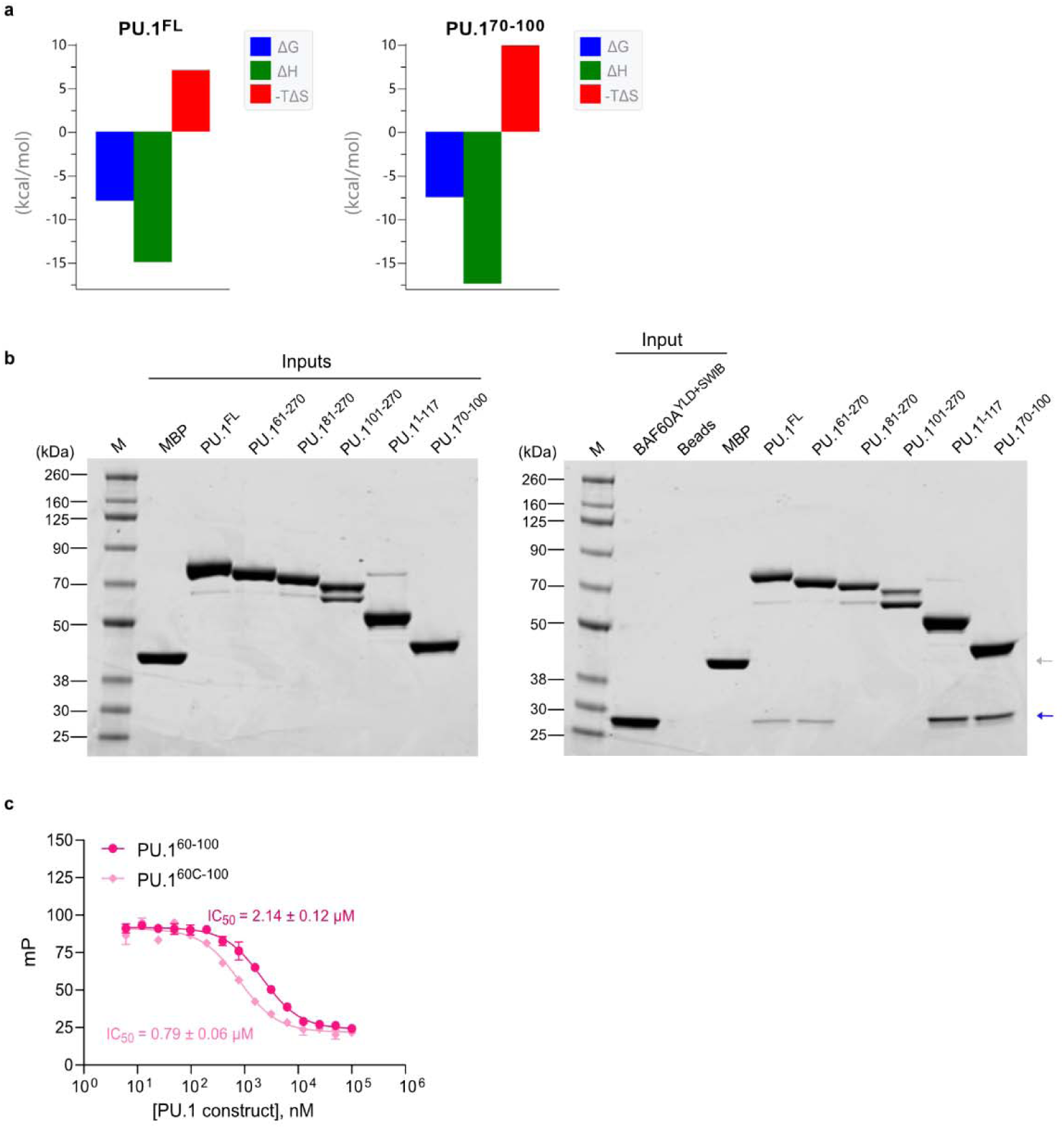
**a,** ITC signature plot of PU.1^FL^ ( left) and PU.1^70–100^ (right) upon binding to BAF60A^YLD+SWIB^. Amylose pull-down assay with PU.1^FL^ and PU.1 truncation constructs demonstrating PU.1^70–100^ is sufficient to bind BAF60A^YLD+SWIB^. Gray arrow indicates MBP only and blue arrow indicates BAF60A^YLD+SWIB^. **c,** FP competition curves for PU.1^60–100^ (darker pink circles) and PU.1^60C-100^ (lighter pink diamonds) competing with PU.1^60–100^^TMR^ for binding to BAF60A^YLD+SWIB^. Errors denote standard deviation from triplicates. Data analyzed in GraphPad Prism (version 10.0.2) using a 4-parameter dose-response curve to obtain IC_50_ ± SEM values.

**Extended Data Fig. 4.**
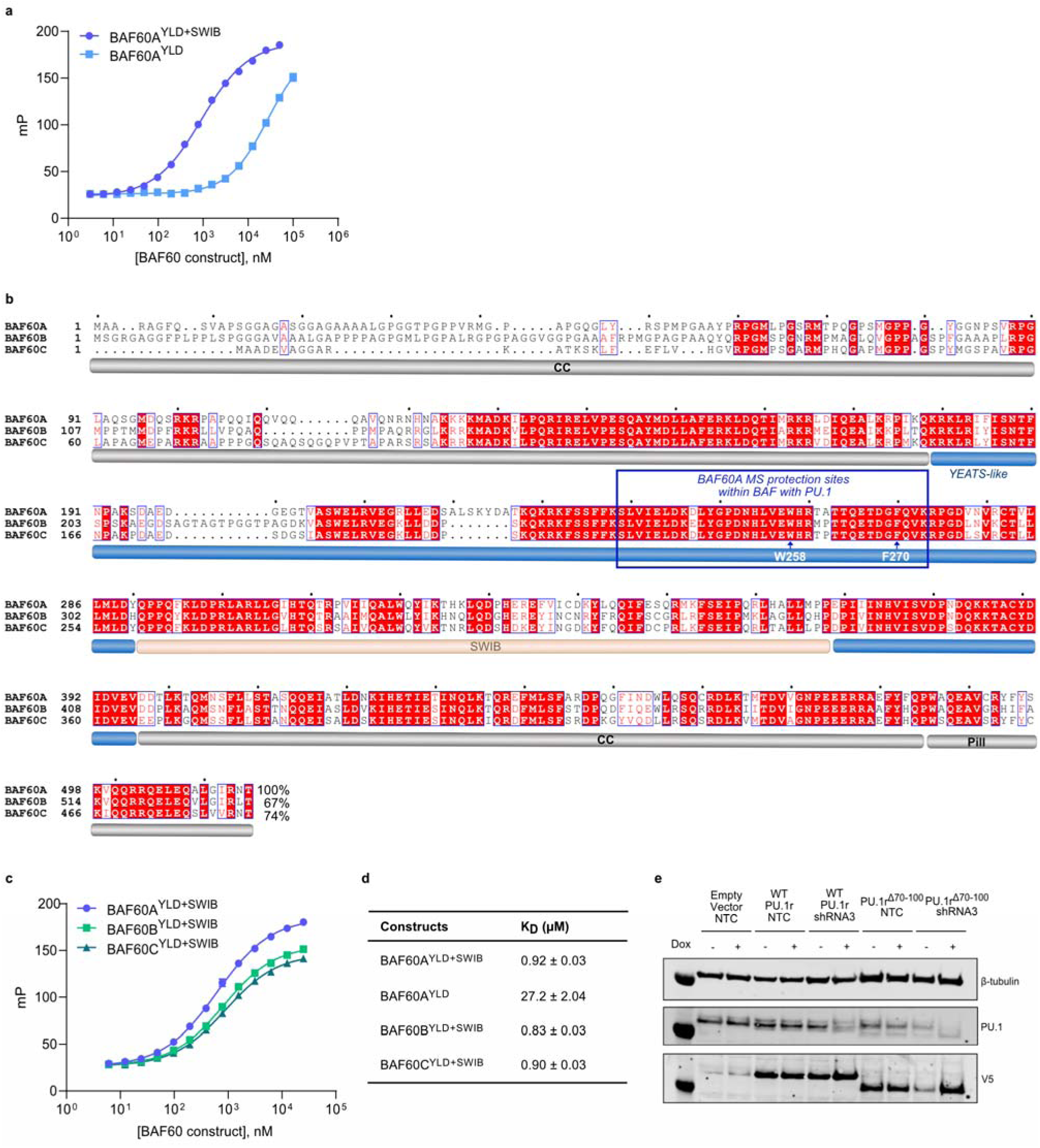
**a,** FP titration curves for binding of PU.1^60–100^^TMR^ to BAF60A^YLD+SWIB^ (blue circles) and BAF60A^YLD^ (blue squares). **b,** Sequence alignment of BAF60 paralogs with domains labeled: coiled coil (CC, dark gray), YLD (blue), SWIB (beige), and the pillar helix (Pill, dark gray). Visualization was generated using the ENDscript server and is colored from low (no box) to high (blue box and red font: similar; red box: identical) sequence identity; percentage sequence identity to BAF60A annotated. The sequence spanning the protected sites identified by MS footprinting is highlighted in a blue box. **c,** FP titration curves for binding of PU.1^60–100^^TMR^ to BAF60A^YLD+SWIB^ (blue circles), BAF60B^YLD+SWIB^ (green squares), and BAF60C^YLD+SWIB^ (green triangles). Errors denote standard deviation from triplicates. **d,** Summary of binding affinities ( K_D_ ± SEM) of PU.1^60–100^^TMR^ to the BAF60 constructs in **a** and **c**. **e,** Western blot showing cellular inputs of MV-4-11 cells containing empty vector, V5-PU.1r, or V5-PU.1r^Δ^^70–100^ with either shPU.1#3 or shNTC (+/- doxycycline).

**Extended Data Table 2.** IC_50_ values of PU.1 mutant and WT PU.1 competing with PU.1^60–100^^TMR^ for binding to BAF60A^YLD+SWIB^ in FP competition assay. Data analyzed in GraphPad Prism (version 10.0.2) using a 4-parameter dose-response curve to obtain IC_50_ ± SEM values.

| Construct | IC<sub>50</sub> (μM) |
| --- | --- |
| WT PU.1 | 0.40 ± 0.08 |
| PU.1<sup>Δ70-100</sup> | >10 |
| PU.1<sup>F70A</sup> | >10 |
| PU.1<sup>L73A</sup> | >10 |
| PU.1<sup>L81A</sup> | >10 |
| PU.1<sup>L84A</sup> | >10 |
| PU.1<sup>D97A</sup> | >10 |
| PU.1<sup>S75A</sup> | 1.71 ± 0.17 |

**Extended Data Fig. 5.**
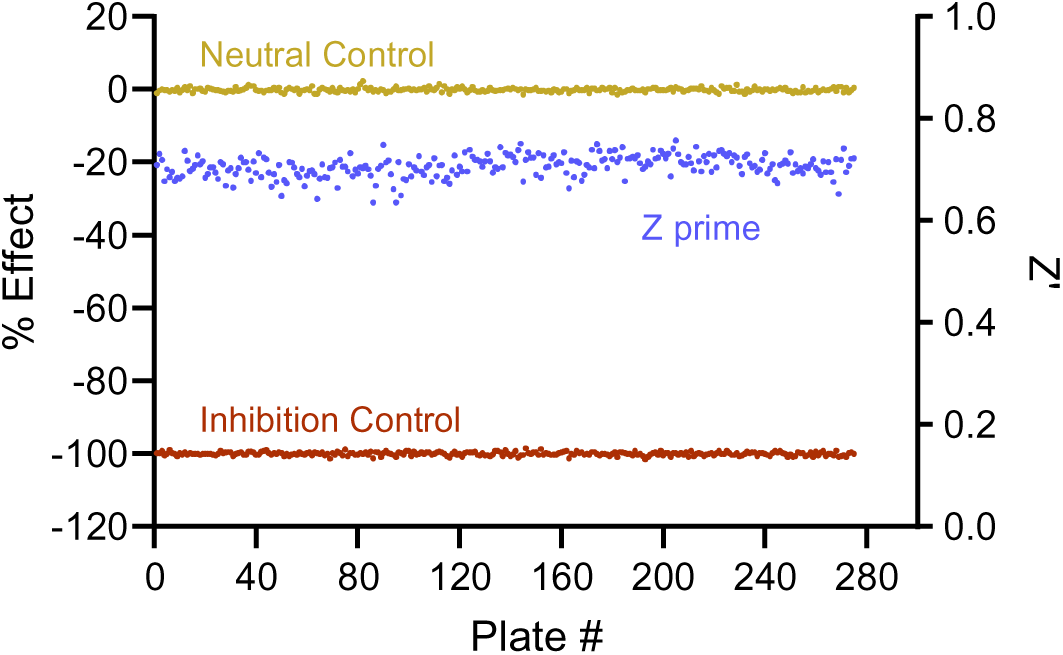
Plot showing screening performance across 280 plates. Neutral control (yellow) and inhibition control (red) are plotted on the left y-axis, while Z’ values (blue) are plotted on the right y-axis. A Z’ value between 0.5 and 1 indicates a high-quality assay well suited to high-throughput screening.

**Extended Data Fig. 6.**
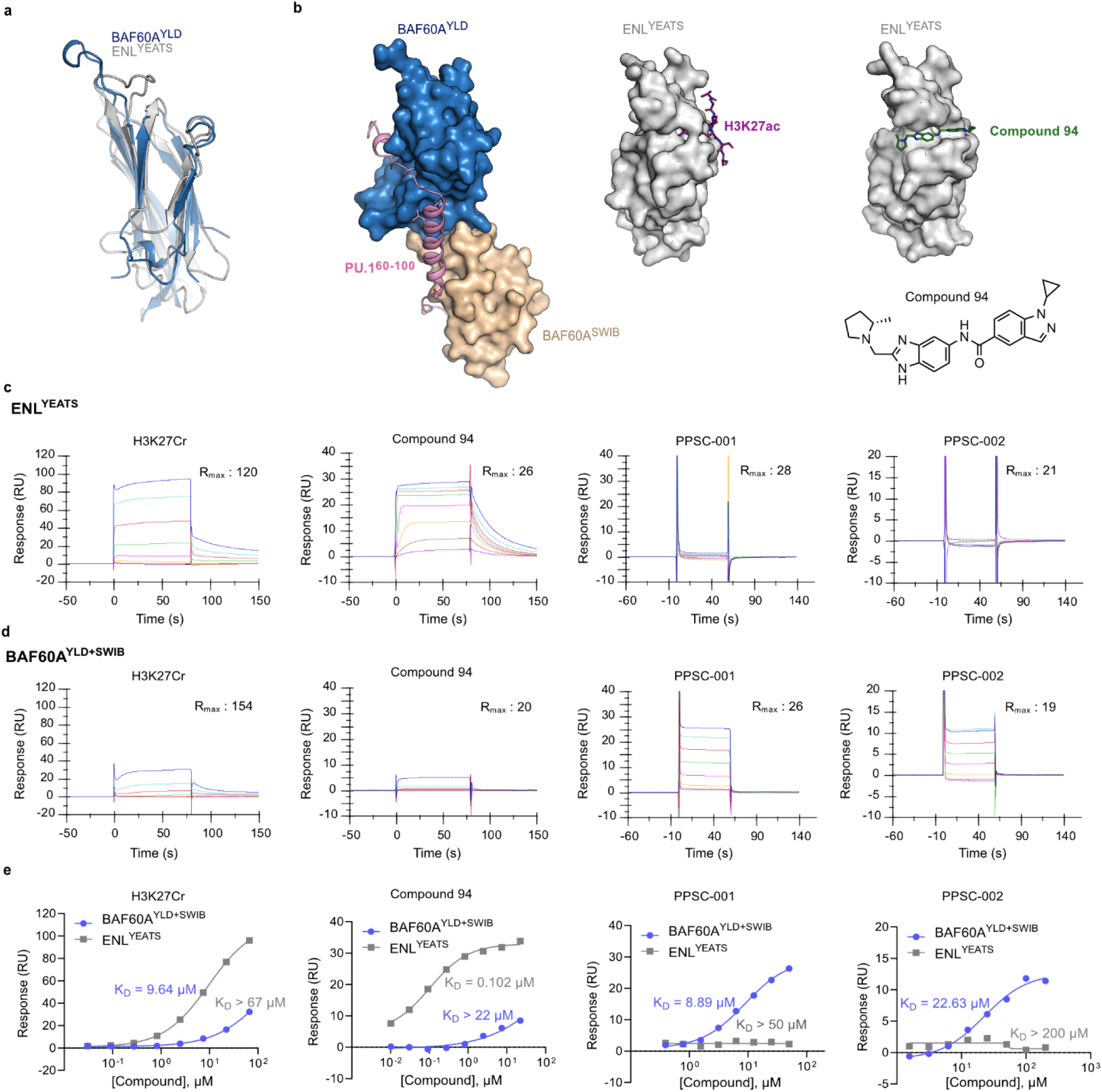
**a,** Superposition of *apo* BAF60A^YLD^ and canonical ENL^YEATS^ (PDB code: 5J9S)^59^. Alignment of 106 Cα residues resulted in a RMSD of 2.19 Å using Coot v0.9.6. **b,** Side-by-side comparison of BAF60A^YLD+SWIB^: PU.1 peptide complex structure (BAF60A^YLD^, surface blue; BAF60A^SWIB^, surface beige; PU.1 peptide, pink cartoon) and ENL^YEATS^ (surface gray) bound with histone H3K27Ac^59^ peptide (purple sticks) and compound 94 (PDB code: 6HT0, green sticks). View from the same orientation shows the histone and PU.1 peptide binding sites are at separate locations in the YEATS fold. Chemical structure of compound 94 that targets YEATS family members, such as ENL^YEATS^ is shown. **c,d,** SPR binding curves for H3K27Cr peptide, Compound 94, PPSC-001, and PPSC-002 binding to **c,** ENL^YEATS^ and **d,** BAF60A^YLD+SWIB^. Names of analytes along with expected saturation response (R_max_) for 1:1 stoichiometry are indicated. **e,** Steady-state affinity measurements of H3K27Cr peptide, Compound 94, PPSC-001, and PPSC-002 binding to ENL^YEATS^ and BAF60A^YLD+SWIB^ by SPR. Equilibrium fits generated in GraphPad Prism (version 10.0.2) using a 4-parameter dose-response curve to obtain K_D_ ± SEM.

